# Rewriting endogenous human transcripts with *trans*-splicing

**DOI:** 10.1101/2024.01.29.577779

**Authors:** Sita S. Chandrasekaran, Cyrus Tau, Matthew Nemeth, April Pawluk, Silvana Konermann, Patrick D. Hsu

**Affiliations:** Arc Institute, 3181 Porter Drive, Palo Alto, CA 94304, USA; Department of Bioengineering, University of California, Berkeley, Berkeley, CA, USA; University of California, Berkeley - University of California, San Francisco Graduate Program in Bioengineering, Berkeley, CA, USA; Center for Computational Biology, University of California, Berkeley, Berkeley, CA, USA; Department of Biochemistry, Stanford University School of Medicine, Stanford, CA, USA

## Abstract

Splicing bridges the gap between static DNA sequence and the diverse and dynamic set of protein products that execute a gene’s biological functions. While exon skipping technologies enable influence over splice site selection, many desired perturbations to the transcriptome require replacement or addition of exogenous exons to target mRNAs: for example, to replace disease-causing exons, repair truncated proteins, or engineer protein fusions. Here, we report the development of <u>R</u>NA-guid<u>e</u>d *trans*-<u>spli</u>cing with <u>C</u>as <u>e</u>ditor (RESPLICE), inspired by the rare, natural process of trans-splicing that joins exons from two distinct primary transcripts. RESPLICE uses two orthogonal RNA-targeting CRISPR effectors to co-localize a trans-splicing pre-mRNA and to inhibit the cis-splicing reaction, respectively. We demonstrate efficient, specific, and programmable trans-splicing of multi-kilobase RNA cargo into nine endogenous transcripts across two human cell types, achieving up to 45% trans-splicing efficiency in bulk, or 90% when sorting for high effector expression. Our results present RESPLICE as a new mode of RNA editing for fine-tuned and transient control of cellular programs without permanent alterations to the genetic code.

## INTRODUCTION

Splicing is a fundamental biological process that plays an important regulatory role at the midpoint of the central dogma of molecular biology, enabling the same genetic sequence to give rise to a dynamic set of protein isoforms that can have distinct biological functions, subcellular localizations, or stabilities (Marasco & Kornblihtt, 2023; Rogalska et al., 2023; Scotti & Swanson, 2016). Errors in splicing regulation are heavily implicated in many disease contexts, including neurodegenerative disorders and muscular dystrophy (Nikom & Zheng, 2023; Pistoni et al., 2010; Scotti & Swanson, 2016). However, our ability to perturb endogenous splicing processes is currently limited to knockdown of specific transcript isoforms or exon-skipping technologies that use antisense oligonucleotides (ASOs) or RNA-targeting CRISPR-Cas systems to influence splice site choice within a given primary transcript (Du et al., 2020; Konermann et al., 2018; Pramono et al., 1996).

The concept of *trans*-splicing, wherein the endogenous splicing machinery is harnessed to join together exons from two different pre-mRNA transcripts, has the potential to provide a broadly applicable platform to manipulate splicing through programmable installation of exon(s) of interest into native transcripts. Efficient and programmable *trans-*splicing technologies could enable the study of oncogenic fusions, install tags or localization sequences, replace exons harboring pathogenic mutations, or repair truncated protein products – all in a transient, reversible manner without permanent alterations to the genetic code.

*Trans*-splicing occurs naturally in rare cases, for instance, joining a short splice leader sequence to mRNAs in trypanosomes and nematodes (Krause & Hirsh, 1987; Ploeg et al., 1982). Human endometrial stromal cells constitutively *trans*-splice exons from *JAZF1* and *JJAZ1* (Li et al., 2008), while a subset of non-small cell lung cancers *trans*-splice *PJA2* and *FER* to generate a chimeric transcript correlated with poor prognosis (Kawakami et al., 2013). Previous studies over the past two decades have attempted to harness this rare *trans*-splicing mechanism by designing an exogenous RNA to both hybridize to a target transcript and supply a *trans*-splicing cargo. This RNA-only system relied on 15-500 bp of RNA-RNA hybridization to localize the *trans*-splicing cargo to the target pre-mRNA and reported *trans-*splicing into highly expressed reporter mini-genes and limited endogenous loci (Puttaraju et al., 1999; Wally et al., 2012). However, these proof-of-concept studies had low efficiency and lacked characterization of specificity, especially for endogenous human transcripts.

Here, we developed an efficient, specific, and programmable *trans-*splicing technology called RESPLICE (<u>R</u>NA-guid<u>e</u>d *trans*-<u>spli</u>cing with <u>C</u>as <u>e</u>ditor) that capitalizes on two key principles: 1) proximity between splice sites is a major determinant of successful splicing reactions, and 2) different splice donors or acceptors exist in competition with each other (Roca et al., 2013; Viles & Sullenger, 2008). RESPLICE employs two modular CRISPR-Cas effectors to take advantage of each of these key principles: one to co-localize the target and cargo pre-mRNA transcript through CRISPR-guided programmable RNA binding, and the other to interfere with the cognate *cis*-splicing reaction through target-specific RNA cleavage downstream of the desired splicing reaction.

We demonstrate *trans*-splicing into 9 endogenous human transcripts across two different cell types, achieving up to 45% *trans*-splicing in unsorted cells and 90% *trans*-splicing in cells sorted for high effector expression with cargo lengths ranging from 800 nt to 2.1 kb. Our transcriptome-wide characterization of *trans*-splicing specificity indicates that off-target capture of the trans-splicing donor can occur across diverse endogenous transcripts, highlighting the promiscuous nature of the splicing machinery. However, apart from the targeted transcript, the amount of off-target trans-splicing was very low on a per-transcript basis, indicating minimal impact on the transcriptome as a whole. We further provide proof-of-concept for therapeutically-relevant *trans-*splicing into pathogenic transcripts with high mutational heterogeneity or repeat expansions. Taken together, our work advances transient, reversible, and large-scale transcriptome engineering for applications across biotechnology and genetic medicine.

## RESULTS

### RNA-targeting CRISPR-Cas13d guides site-specific *trans*-splicing

In an attempt to overcome the limitations of prior *trans*-splicing technologies that utilized an RNA-only approach, we first designed a CRISPR-guided *trans*-splicing system using our previously reported RNA-targeting Cas13d from *Ruminococcus flavefaciens* XPD3002 (CasRx) (Konermann et al., 2018). We paired the catalytically inactive form of the enzyme, dCasRx, with a guided *trans-*splicing module (TSM): an RNA consisting of 1) a guide containing a CasRx direct repeat (DR) and a spacer with reverse complementarity to the target sequence (Konermann et al., 2018); 2) a minimal intron containing a stuffer region and splice sites for spliceosomal recognition (Uckun et al., 2015); and 3) a cargo sequence containing the desired exon to be *trans*-spliced (**Fig. 1B**). We sought to test the ability of dCasRx to enhance the efficiency and specificity of the system by targeting the TSM to the target pre-mRNA in a guide RNA-programmable manner.

**Figure 1:**
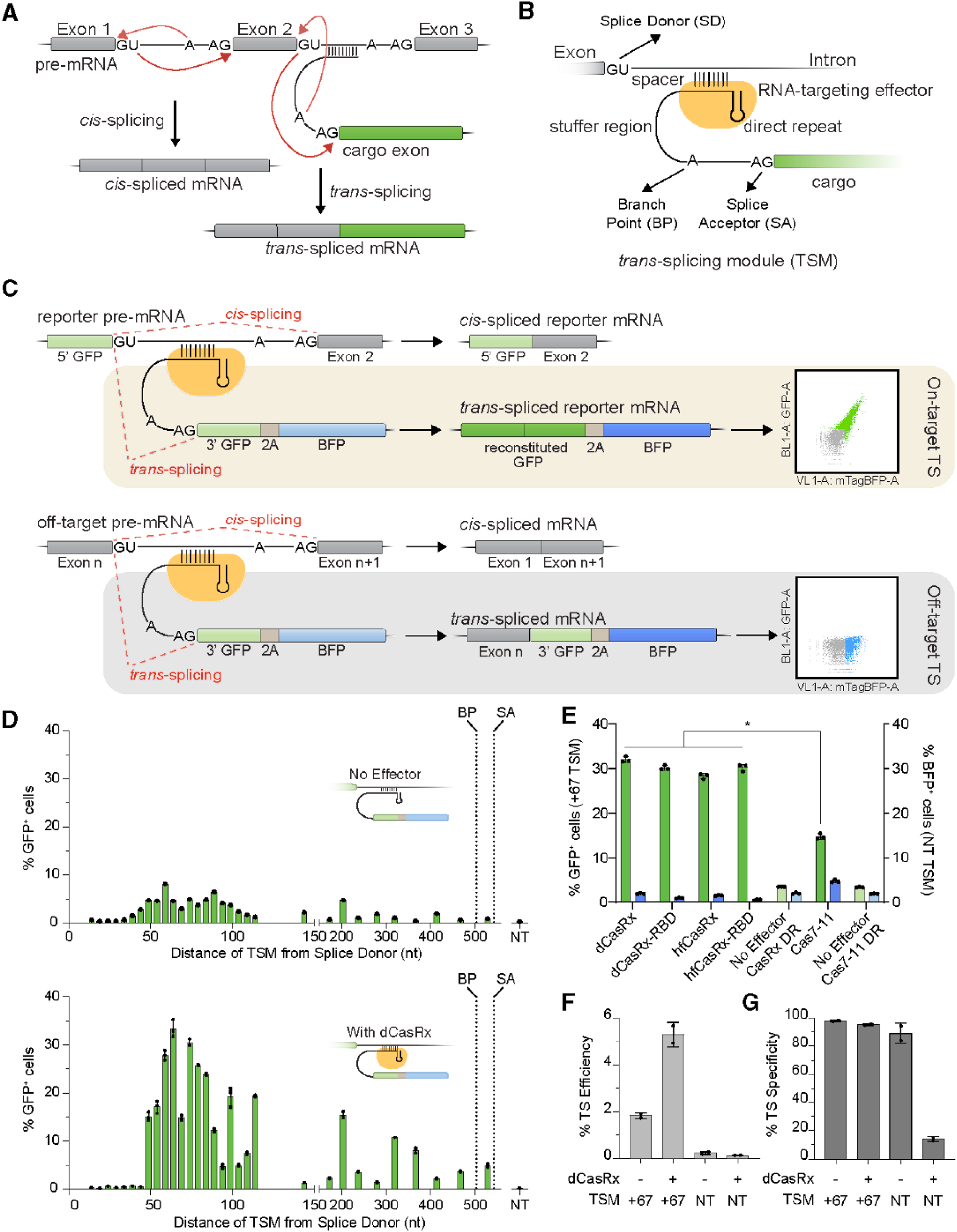
An RNA-targeting CRISPR effector guides site-specific *trans*-splicing. **(a)** Schematic representation of the *cis*- and *trans*-splicing reactions. Exons are denoted by boxes, lines denote introns, splice donors are represented by ’GU’, branch points (BP) are represented by ’A’, and splice acceptors (SA) are represented by ’AG’. Green box represents the exon splicing *in trans*. **(b)** Design of the guided *trans*-splicing module (TSM), including an RNA-targeting CRISPR-Cas effector presenting a guide RNA spacer that can hybridize to the target pre-mRNA, a stuffer region, a splice region containing splice branch point and acceptor sites, and the cargo exon(s) to be *trans*-spliced. **(c)** The bichromatic *trans*-splicing reporter enables the assessment of relative levels of on- and off-target *trans*-splicing (TS) events. The reporter pre-mRNA transcript consists of the 5’ half of the GFP coding sequence (CDS), followed by an intron and a second exon. *Cis*-splicing of this reporter results in a truncated GFP protein that does not fluoresce. However, *trans*-splicing of the TSM carrying the 3’ half of the GFP CDS as a cargo exon with the reporter pre-mRNA reconstitutes the full length GFP CDS, followed by a 2A ribosomal skip sequence and full-length BFP, yielding cells that express both GFP and BFP. Any off-target *trans*-splicing of the TSM into non-reporter pre-mRNAs would only result in the expression of BFP – provided that the reaction placed BFP in-frame with the upstream coding sequence due to lack of a start codon. **(d)** Application of the bichromatic reporter in HEK293T cells to assess efficiency of *trans-*splicing, assessed by % GFP⁺ cells, with (bottom) or without (top) the presence of the TSM effector, dCasRx. Efficiency was compared across 29 TSM spacers tiling the reporter intron, with locations indicated on the x-axis. The locations of the branch point (BP) and splice acceptor site (SA) are labeled for reference. A TSM encoding a non-targeting spacer (NT) was included as a negative control, shown on the right. Data are mean ± s.d.; n = 3 biological replicates; individual data points are shown. **(e)** A comparison of efficiency and specificity across TSM effectors listed on the x-axis (RNA-binding domain, RBD; HiFiCasRx, hfCasRx; direct repeat, DR). Percentage of single GFP^+^ HEK293T cells– as a readout of on-target *trans-*splicing efficiency – is plotted on the left y-axis in green bars for the TSM spacer targeting 67 nucleotides downstream of the splice donor on the reporter transcript. Percentage of single BFP^+^ HEK293T cells is plotted on the right y-axis in blue bars for the non-targeting (NT) TSM spacer, as a measure of relative off-target levels. Significance testing between Cas7-11 and dCasRx was performed with an unpaired two-tailed Welch’s test assuming unequal variances. Data are mean ± s.d.; n = 3 biological replicates; individual data points are shown. **(f)** Percentage of on-target *trans*-splicing efficiency into the reporter mRNA in HEK293T cells as measured by RNA-seq (*trans-*spliced / total reporter junctions) with or without the dCasRx effector and using either the +67 nt TSM spacer or a non-targeting (NT) spacer. Data are represented by mean ± s.d.; n = 2 biological replicates, with individual points plotted. **(g)** Percentages of on-target *trans*-splicing specificity measured by RNA-seq of HEK293T cells, defined as the number of on-target junction reads divided by the total number of transcriptome-wide *trans*-splicing junctions passing filter (see **Methods**), for reactions with or without the dCasRx effector and using either the +67 nt TSM spacer or a non-targeting (NT) spacer. Data are represented as mean ± s.d.; n = 2 biological replicates, with individual points plotted.

In order to assess the ability of this system to direct target-specific *trans*-splicing in HEK293T cells, we next designed a plasmid-encoded reporter transcript by combining the N-terminal half of GFP with an intron and exon derived from a previously established alternative splicing reporter (Orengo et al., 2006) (**Fig. 1C**). We targeted the reporter intron with TSMs carrying a 1.1kb cargo consisting of the C-terminal half of GFP followed by full-length BFP. Successful on-target *trans*-splicing of the reporter would therefore result in the reconstitution of an intact GFP sequence, yielding GFP⁺/BFP⁺ cells that can be quantified by flow cytometry, whereas off-target *trans*-splicing into an in-frame ORF would result in the production of BFP only (**Fig. 1C**). We designed TSMs to tile the reporter intron with 26 different guide sequences, or a non-targeting (NT) guide, while keeping the rest of the TSM constant. HEK293T cells were cotransfected with each TSM alongside the reporter plasmid, and either the dCasRx expression plasmid or pUC19 as an effector-less negative control. Using flow cytometry, we then quantified the percentage of single cells expressing GFP and BFP.

Consistent with previous RNA-only *trans*-splicing attempts, we found that supplying the TSM only without an effector yielded up to 5% GFP⁺ cells (**Fig. 1D**). Strikingly, the dCasRx effector enabled a consistent ∼6-fold improvement of *trans*-splicing efficiency compared to the no-effector control: up to 30% (**Fig. 1D**). We noted a peak in *trans*-splicing efficiency when the TSM was targeted to the region ∼50-100 nt downstream from the splice donor site, where we could typically detect ∼30% GFP⁺ cells (**Fig. 1D**).

Having established that an RNA-binding CRISPR effector can substantially improve the efficiency of *trans*-splicing, we next aimed to assess a panel of different RNA-targeting effectors to optimize our system. We tested hfCasRx, due to its reduced collateral and off-target cleavage (Tong et al., 2022), dCasRx and hfCasRx with dsRBD fusions, due to the fusion’s improved specificity with dCasRx in RNA imaging applications (dCasRx-RBD and hfCasRx-RBD) (Han et al., 2020), and DiCas7-11 (Cas7-11), also due to its reportedly low collateral cleavage and its orthogonality to the Cas13 family (Özcan et al., 2021) – each with cognate direct repeat (DR) sequences in their corresponding TSM constructs. For all effectors, the +67 nt targeting location produced the highest efficiency of on-target *trans*-splicing as indicated by the percentage of GFP⁺ cells (**Fig. 1E, Fig. S1A**). hfCasRx, dCasRx-RBD, and hfCasRx-RBD had slightly lower *trans*-splicing efficiencies than dCasRx. However, these effectors also exhibited 2-3 fold decreased BFP expression, suggesting a potential trade-off of on-target efficiency for higher specificity (**Fig. 1E**). On the other hand, Cas7-11 had only half the percentage of GFP⁺ cells while exhibiting 2-fold higher BFP⁺ cells compared to dCasRx – suggesting lower efficiency and specificity for this effector, in agreement with lower levels of transcript knockdown and higher off-target effects of Cas7-11 reported previously (Wei et al., 2023; Zeballos C. et al., 2023). These general trends for effector performance held true across three additional target sites: one located further downstream within the intron (+470 nt), one targeting the splice acceptor sequence (+531 nt), and one in the second exon (+546 nt) (**Fig. S1A**). Therefore, we chose the dCasRx effector, which showed the highest *trans*-splicing activity, to continue developing and characterizing the system. For certain applications requiring higher specificity or RNA cleavage at the TSM targeting site, the dsRBD fusions or hfCasRx may be advantageous.

In our reporter construct, we also noticed strong promoter-dependent variability in the percentages of GFP⁺ and BFP⁺ cells using the same TSM and effector combination (**Fig. S2A**). This was reminiscent of previous work showing that some promoters may have mechanistically independent and difficult-to-interpret effects on RNA editing tools (Kaseniit et al., 2022). To mitigate this, we chose to use the SFFV promoter due to its highest GFP⁺/BFP⁺ signal-to-noise ratio (**Fig. S2B**), in agreement with prior work suggesting SFFV minimizes these potential artifacts (Kaseniit et al., 2022).

A major challenge in previous efforts to develop *trans*-splicing systems has been the inability of most experimental setups to profile transcriptome-wide off-target events. In fact, the most comprehensive off-target analyses of *trans*-splicing in the literature performed “one-sided” RT-PCR of the *trans*-splicing cargo and either visualized these products on a gel (Puttaraju et al., 1999) or sequenced clones, in which case all twenty analyzed clones represented off-target – not on-target – *trans*-splicing events (Kikumori et al., 2001). The bichromatic *trans*-splicing reporter readout we used in the above experiments similarly does not allow us to comprehensively detect or quantify transcriptome-wide off-targets of our system: its protein-based fluorescent readout does not directly represent RNA-level changes, BFP would not be expressed from out-of-frame off-targets, and basal BFP expression from the un-spliced TSM would confound any accurate quantification.

To more accurately quantify the on- and off-target performance of our dCasRx-based *trans*-splicing system at the transcriptome-wide level, we performed RNA-seq and compared the +67 site against the NT guide with an effector-less negative control. We calculated *trans*-splicing efficiency as a ratio of *trans*-spliced junction reads to the sum of *trans* and *cis* splice junction reads, measuring an efficiency of 5.3% in the presence of dCasRx: a five-fold increase from the corresponding no-effector condition (**Fig. 1F)**. Off-targets with dCasRx were low, with only an average of 137 normalized off-target *trans*-splicing junction reads across the two replicates, compared to an average of 2837 normalized on-target *trans*-splicing junction reads, leading to an average on-target specificity of 95.3% (**Fig. 1G**, **Table S1**). Of these, only 7 out of 62 total off-target *trans*-splicing junctions were commonly detected across replicates (**Table S2**). Notably, all of the off-target *trans*-splicing junctions were represented by fewer than ten out of a total 145 million reads, indicating that these aberrant splice products exhibited very low abundance (**Methods**). Additionally, over half of the off-target loci had less than 5% *trans*-splicing efficiency, indicating that these transcripts were *cis-*spliced in the majority of cases, maintaining the expression of unperturbed transcript isoforms. Interestingly, we also discovered two additional on-target splice isoforms for the reporter, which likely result from an artifact of the alternative splicing reporter (**Fig. S3**). While we observed *trans*-splicing into these different reporter splice donors, successful alternate *trans*-splicing would not result in the production of full-length GFP (**Fig. S3**).

We conclude that, while we can detect off-target *trans*-splicing at multiple loci, each of these off-targets occurs at low efficiency and with substantial stochasticity. Taken together, our proof-of-concept experiments demonstrate that CRISPR-guided *trans*-splicing can direct programmable, site-specific, and efficient RNA writing in HEK293T cells.

### Improving CRISPR-guided *trans*-splicing efficiency by inhibiting *cis*-splicing with a second CRISPR ribonuclease

Any *trans*-splicing reaction must compete with the endogenous *cis*-splicing reaction that would pair the target exon with its downstream cognate exon (Viles & Sullenger, 2008). In our RNA-seq analysis, we observed that over 94% of the reporter construct underwent *cis*-splicing, even with our most effective *trans*-splicing system. In an attempt to shift the balance toward the desired *trans*-splicing reaction, we took inspiration from previous work using ASOs to inhibit the usage of specific splice acceptors (Coady et al., 2008; Liemberger et al., 2018). Instead of an ASO, we added a second, catalytically-active CRISPR effector, which we term the *cis* interfering module (CIM) (**Fig. 2B**), to cleave the target transcript downstream of the TSM binding site (**Fig. 2A**). We tested 3 different CIM effectors (Cas7-11, hfCasRx, and hfCasRx-RBD) targeting 3 different locations (+210, +492, and +531 nt) in the reporter intron and paired them with the TSM effector and targeting location showing the highest on-target efficiency (dCasRx and +67 TSM).

**Figure 2:**
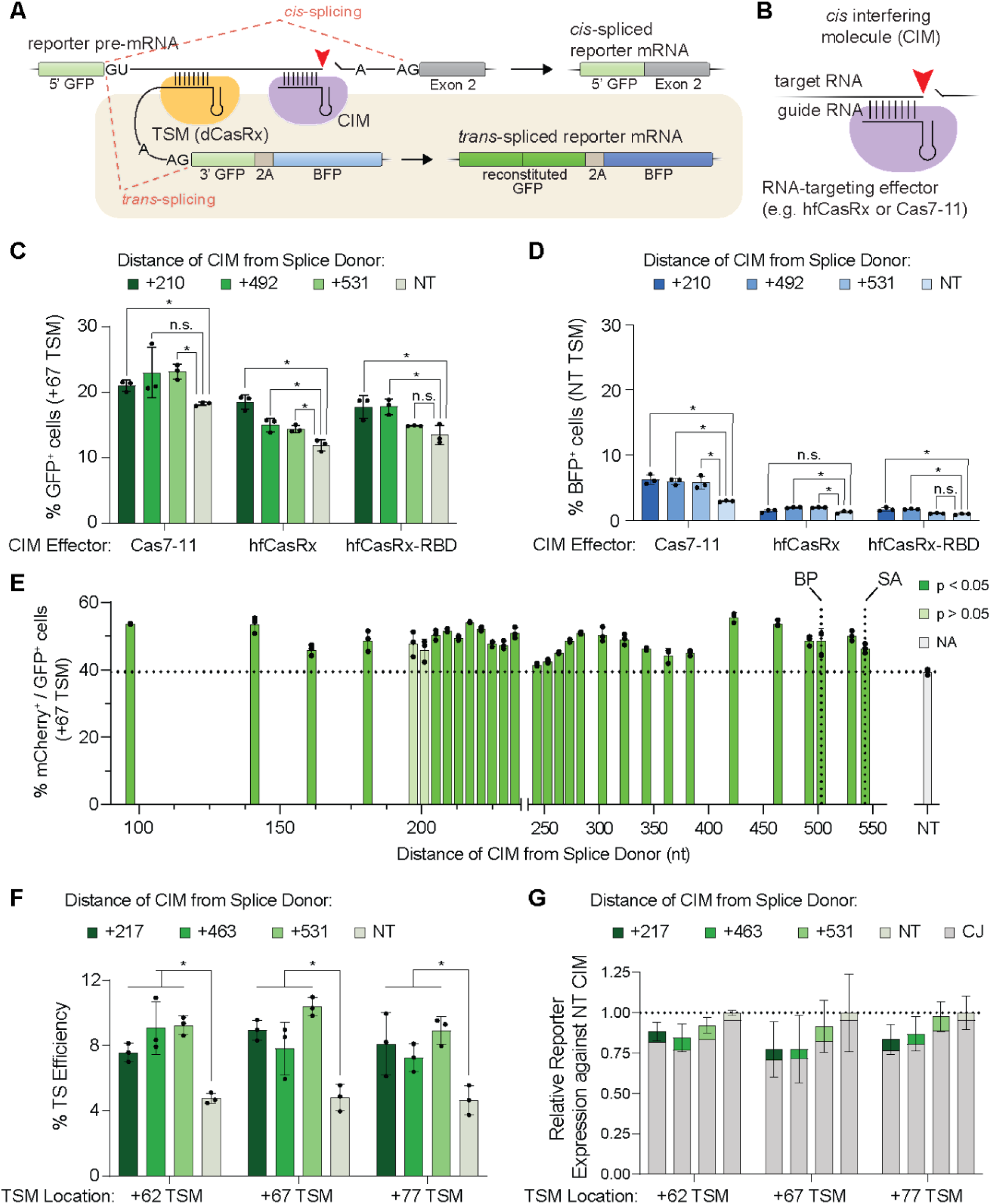
A *cis-splicing* interfering module (CIM) boosts *trans-*splicing efficiency. **(a)** Schematic of the bichromatic *trans*-splicing reporter system, as in Figure 1C, with the addition of a *cis* interfering module (CIM; purple). The catalytically active CIM effector is designed to cleave the reporter transcript at its target site, downstream of where the TSM effector (yellow) binds. **(b)** Schematic of the CIM. The CIM consists of a catalytically active RNA-guided, RNA-targeting CRISPR effector (Cas7-11, hfCasRx, or hfCasRx-RBD) bound to its cognate guide RNA, which will direct site-specific target transcript cleavage to inhibit the *cis-*splicing reaction. **(c)** On-target reporter *trans*-splicing efficiency, measured by the percentage of single GFP^+^ HEK293T cells, of the dCasRx-bound +67 TSM paired with either Cas7-11, hfCasRx, or hfCasRx-RBD as the CIM effector. We tested three targeting CIM spacers (+210 nt, +492 nt, +531 nt) or a non-targeting CIM guide (NT). Significance between targeting and non-targeting spacers were performed with an unpaired two-tailed Welch’s t-test assuming unequal variances. Data are represented as mean ± s.d.; n = 3 biological replicates, with individual points plotted. **(d)** Comparison of relative specificity across different CIM effectors and spacer targeting locations, as measured by single BFP⁺ HEK293T cells, using a dCasRx-bound NT TSM paired with each indicated CIM effector. We tested three targeting CIM spacers (+210 nt, +492 nt, +531 nt) or a non-targeting CIM guide (NT). Significance between targeting and non-targeting spacers were performed with an unpaired two-tailed Welch’s test assuming unequal variances. Data are represented as mean ± s.d.; n = 3 biological replicates, with individual points plotted. **(e)** Comparison of % single GFP⁺ HEK293T cells indicating *trans-*splicing efficiency the +67 TSM for each tested CIM tiling the reporter. Significance between targeting and non-targeting spacers were performed with an unpaired two-tailed Welch’s t-test assuming unequal variances. Due to the high number of comparisons, we corrected p-values using the Benjamini Hochberg multiple testing procedure. Targeting locations that were not significantly different from the non-targeting CIM are indicated with lighter bars. Data are mean ± s.d.; n = 3 biological replicates, with individual points plotted. **(f)** On-target reporter *trans*-splicing efficiency, measured by ddPCR, of a dCasRx-bound TSM paired with a Cas7-11 CIM effector in HEK293T cells. We tested three targeting CIM spacers (+217 nt, +463 nt, +531 nt) or a non-targeting CIM guide (NT) paired with three targeting TSM spacers (+62 nt,+67 nt,+77 nt). Significance between targeting and non-targeting spacers were performed with an unpaired two-tailed Welch’s test assuming unequal variances. Data are represented as mean ± s.d.; n = 3 biological replicates, with individual points plotted. **(g)** Fold change of reporter expression averaged across three replicates of each Cas7-11 CIM (+217 nt, +463 nt, +531 nt, NT relative to the NT CIM for all the TSM spacers shown in panel F (+62 nt,+67 nt,+77 nt) measured by ddPCR in HEK293T cells. The gray portion of the bar corresponds to the proportion of *cis* junctions, while the green portion of the bars correspond to the proportion of *trans* junctions and indicates the CIM position used.

We found that we could consistently increase the percentage of GFP⁺ cells by approximately 1.2-fold upon providing a targeting Cas7-11 CIM (**Fig. 2C**). While hfCasRx and hfCasRx-RBD had higher fold increases of GFP⁺ cells (1.6-fold and 1.3-fold for +210 nt CIM spacer relative to the NT CIM spacer), they had lower absolute efficiency. We hypothesize this could be due to the high expression of the U6 promoter-driven CIM recruiting dCasRx away from the TSM DR, since the hfCasRx guide sequences share the same dCasRx DR. However, consistent to what we observed with the TSM effector comparison, provision of a Cas7-11 CIM increases BFP expression, our proxy for in-frame, off-target *trans*-splicing events, even in non-targeting TSM conditions, suggesting that the promiscuity of Cas7-11 off-target activity could lead to greater off-target *trans*-splicing (Wei et al., 2023; Zeballos C. et al., 2023) (**Fig. 2D**). Choosing here to maximize on-target *trans*-splicing efficiency, we decided to move forward with the Cas7-11 CIM effector despite this potential limitation. We targeted CIMs to 31 different sites between 100-575 nucleotides downstream of the splice donor. Among cells receiving the TSM and CIM effectors, we detected up to 55% GFP⁺ cells, representing up to a 1.4-fold increase in efficiency over a non-targeting CIM control (**Fig. 2E**).

To further validate our results that indicate CIM’s ability to boost trans-splicing, we performed digital droplet PCR (ddPCR) to quantify the *trans*-spliced and *cis*-spliced reporter junctions for the top performing spacers. We chose the top 3 performing TSM spacers (+62, +67, +77) and combined them with each of 3 high performing CIM spacers throughout the reporter intron (+217, +463, +531) along with non-targeting controls. We observed an approximate 2.0-fold increase in *trans*-splicing efficiency between the non-targeting and +531 CIM spacers (from ∼8% to ∼16%) for each of the TSM-CIM pairs tested in the presence of dCasRx (**Fig. 2F**). Furthermore, the rank-order of CIM performance was reasonably consistent across the different TSM spacers, both with and without TSM effectors, implying that CIMs work orthogonally to TSMs (**Fig. 2F, Fig. S4B**). Quantitative analysis of *cis* and *trans* junction reads confirmed that total expression level of the target transcript was within 25% of the corresponding non-targeting CIM condition across all conditions, indicating only mild knockdown by the CIM effector, whereas an increase in *trans* junction reads was the primary driver of doubled *trans-*splicing efficiency (**Fig. 2G**). We termed the combination of the TSM and CIM to drive efficient *trans*-splicing RESPLICE (<u>R</u>NA-guid<u>e</u>d *trans*-<u>spli</u>cing with <u>C</u>as <u>e</u>ditor).

### Efficient on-target *trans*-splicing of endogenous human transcripts

Previous attempts at *trans*-splicing reported in the literature achieved success with highly-expressed and/or simplified (i.e. minigene) reporter constructs but did not comprehensively or directly characterize efficiency or transcriptome-wide specificity when targeting endogenous transcripts (Kikumori et al., 2001; Liu et al., 2002; Mansfield et al., 2000; Murauer et al., 2011; Puttaraju et al., 1999; Tockner et al., 2016; Uckun et al., 2015). Since endogenous targets are at the core of most potential applications of *trans*-splicing for research and medicine, we sought to move RESPLICE beyond an artificial reporter system to characterize its ability to *trans*-splice into endogenous human transcripts at the molecular level. In order to do this, we designed a 791 nt TSM cargo encoding in-frame, full-length msfGFP and first explored on-target efficiency using a ddPCR readout to measure the ratio of *trans*-spliced junctions to *cis*-spliced junctions (**Fig. 3A)**.

**Figure 3:**
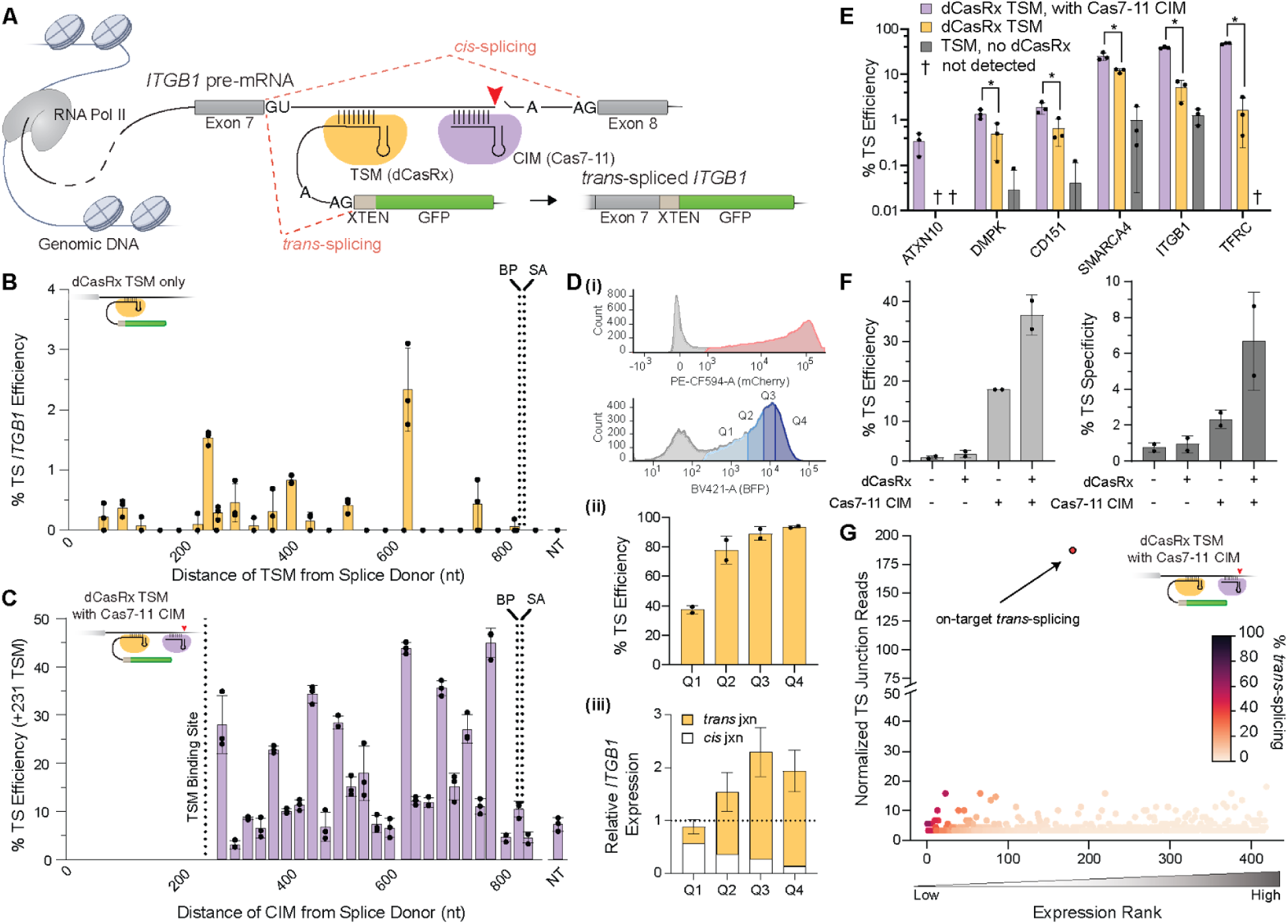
RESPLICE *trans*-splices cargo exons onto endogenous human transcripts. **(a)** Schematic depicting *trans*-splicing into an endogenous pre-mRNA transcript – in this case, *ITGB1* – with a TSM targeting downstream of exon 7 and a CIM targeting downstream of the TSM. Splice donors, branch points, and acceptors are labeled as in Figure 1a. **(b)** *Trans*-splicing efficiency as measured by ddPCR (*trans-*spliced junction reads over total exon 7 junction reads) of 29 TSM spacers targeting sites across intron 7 of *ITGB1*, or non-targeting (NT) TSM with a dCasRx effector in HEK293T cells. Branch point (BP) and splice acceptor (SA) sites are labeled for reference. Data are mean ± s.d.; n = 3 biological replicates, with individual data points shown. **(c)** *Trans*-splicing efficiency into *ITGB1* as measured by ddPCR in HEK293T cells, using a dCasRx-bound TSM targeting 231 nucleotides downstream of the splice donor of intron 7 in ITGB1 together with one of 27 Cas7-11 CIMs targeting downstream of the TSM target site (or NT CIM). Branch point (BP), splice acceptor (SA), and TSM binding site are labeled for reference. Data are mean ± s.d.; n = 3 biological replicates, with individual points plotted. **(d) (i)** Flow cytometry measurements of transfected HEK293T cells expressing mCherry (as a proxy for dCasRx TSM effector expression; top) or mTagBFP2 (indicating Cas7-11 CIM effector expression; bottom) from a single transfection. The red shading in the mCherry plot indicates HEK293T cells that were sorted as positive for dCasRx expression, and the blue shaded quartiles Q1-Q4 indicate the sorting performed to stratify cells by their level of Cas7-11 expression for further analysis. **(ii)** *Trans*-splicing efficiency, as measured by ddPCR, among sorted cell populations scored as positive for dCasRx expression and representing quartiles of Cas7-11 expression (Q1-Q4), using a +231 nt dCasRx TSM in conjunction with a +633 nt Cas7-11 CIM. Data are mean ± s.d.; n = 2 biological replicates, with individual points plotted. **(iii)** The bars represent the average relative proportion of total *ITGB1* transcripts that are *cis*- or *trans*-spliced, as measured by ddPCR, for dCasRx+ cells split into Cas7-11 expression quartiles (Q1-Q4) across two replicates. Each dot represents the relative total expression of *ITGB1* (total number of *cis*- and *trans*-spliced junctions) compared to the average of both replicates of a negative control (NT TSM with NT CIM) in each quartile. **(e)** *Trans*-splicing efficiency across 6 endogenous transcripts in HEK293T cells using the highest efficiency dCasRx TSM and Cas7-11 CIM found through the TSM and CIM screens (**Fig. S5B, S5C**). The negative controls received only an effectorless TSM with no CIM. Data are mean ± s.d.; n = 3 biological replicates, with individual points plotted. **(f)** Efficiency and specificity of the *trans*-splicing of *ITGB1* in HEK293T cells by RNA sequencing. The plot on the left shows the percentage of total *ITGB1* exon 7 junction reads that are *trans*-spliced. The plot on the right indicates the percentage of all transcriptome-wide cargo exon-containing reads that map to the desired on-target *ITGB1* transcript. The TSM targets +231 nt into intron 7, and the CIM consists of a Cas7-11 with a guide targeting +633 nt into intron 7. Data are mean ± s.d.; n = 2 biological replicates, with individual points plotted. **(g)** A snapshot of *trans*-splicing efficiency and specificity across all detected *trans*-splicing (i.e. cargo exon-containing) junctions in HEK293T cells from RNA-seq that passed filtering for the best *ITGB1* TSM and CIM condition (+231 TSM bound to dCasRx and +633 Cas7-11 CIM) for one replicate (other replicate shown in **Fig. S7**). Each dot represents a specific *trans*-splicing junction where the y-axis denotes number of normalized *trans*-splicing junction reads, the x-axis is the transcriptome-wide expression rank of each junction (low expression on the left to high on the right), and the color of each dot corresponds to the *trans*-splicing efficiency of that junction (*trans-*spliced reads divided by total [*trans* + *cis*] junction reads), according to the legend on the right.

We first targeted intron 7 of the *ITGB1* transcript, which encodes integrin subunit beta 1, a highly expressed gene that is not essential for HEK293T viability (Nuñez et al., 2021). We designed TSMs targeting 29 sites downstream of the splice donor (+71 nt to +841 nt) using a Cas13d guide design algorithm to nominate highly efficient guides (Wei et al., 2023) alongside a set of guides to tile the intron at regular intervals. We detected *trans*-splicing when targeting 15 of the 29 sites tested, with efficiencies ranging from 0.2-3%: comparable to the GFP reporter *trans*-splicing efficiencies we measured above by RNA-seq without the addition of a CIM (**Fig. 3B, 1F)**.

Since we found above that the addition of a Cas7-11 CIM increased the efficiency of *trans*-splicing on our reporter, we designed Cas7-11 CIMs to target 27 locations downstream of a high efficiency TSM at +231 nt to reduce *cis-*splicing (**Fig. 3C)**. Strikingly, we found that addition of the CIM boosted the efficiency of *trans*-splicing by more than 20-fold in this endogenous context, reaching 40.1% when targeting the +604 site, as measured by ddPCR (**Fig. 3C)**. We next analyzed the top three performing guides for total ITGB1 transcript abundance and found minimal perturbation of total *ITGB1* transcript amount from the NT CIM control (**Fig. S4A)**.

For both the TSM and CIM, we noticed that efficiency predictions from the nuclease active CasRx guide design algorithm (Wei et al., 2023) were moderately correlated with *trans*-splicing efficiency. In our tiling of *ITGB1* TSMs, we found that the predicted and ddPCR-determined efficiencies had a Pearson correlation of 0.69 (p=1.5E-05) (**Fig. S4B**). Similarly, in our tiling of *ITGB1* CIMs, the predicted and actual efficiencies had a Pearson correlation of 0.64 (p=2.9E-04) (**Fig. S4D**), suggesting that use of this guide design tool can prioritize TSM and CIM targeting sequences most likely to direct high efficiency *trans*-splicing.

Since our primary *trans*-splicing efficiency metric is calculated from unsorted cells, we reasoned that the *trans-*splicing efficiency could be limited by the transfection efficiency of our multi-plasmid system. To better understand the efficiency when all components of the system are present, we sorted cells based on the presence of dCasRx (using mCherry expression as proxy) and binned the dCasRx+ population into quartiles by Cas7-11 expression (as measured by BFP expression) (**Fig. 3D (i, ii)**). We found that dCasRx+ cell populations with high expression levels of Cas7-11 exhibited ∼90% *trans*-splicing of the *ITGB1* target, without reducing total *ITGB1* transcript levels relative to the non-targeting TSM/non-targeting CIM control (**Fig. 3D, Fig. S5A**). Overall, we conclude that the CIM with nuclease active Cas7-11 promotes *trans*-splicing on endogenous human transcripts to high levels, without significantly reducing total target transcript expression. In addition, our data indicates that cell sorting on effector expression can significantly increase *trans*-splicing efficiencies for applications where that might be feasible, yielding very high levels of *trans*-splicing.

Next, we extended this analysis to a set of five additional endogenous target transcripts (*ATXN10*, *DMPK*, *CD151*, *SMARCA4*, and *TFRC*), spanning diverse target intron lengths (330 nt - 54,816 nt), expression levels (21-156 TPM), and functions (**Table S3**). To efficiently identify top-performing TSM-CIM combinations for each target transcript, we used the guide efficiency prediction tool combined with screening (Wei et al., 2023) to identify the top-performing TSM, which we then paired with three top predicted CIM spacers to arrive at the best TSM-CIM combination (**Fig. S5B, S5C**). The best combination for each target transcript resulted in the following on-target *trans*-splicing efficiencies, when measured by ddPCR in unsorted cells: 47.4% for *TFRC*, 17.4% for *SMARCA4*, 2.1% for *DMPK*, 1.9% for *CD151* and 0.3% for *ATXN10*; these efficiencies were consistent with a qPCR readout (**Fig. 3E, Fig. S5B**). Consistent with our data from the splicing reporter and *ITGB1* tiling experiments, we observed an increase in *trans*-splicing efficiency across all endogenous targets by the addition of a CIM to block *cis*-splicing. This increase ranged from 2.8-fold (0.5% to 1.3%) for *DMPK*, to an impressive 28.8-fold (1.6% to 47.4%) for *TFRC* (**Table S3**).

The overall *trans*-splicing efficiency in the presence of both the TSM and CIM was highest for targets with intermediate intron sizes (849 nt for *ITGB1* with 47.4% efficiency, 1,827 nt for *TFRC* with 47.4% efficiency, and 6,180 nt for *SMARCA4* with 17.4% efficiency) (**Table S3**). Given the high efficiency of *trans-*splicing into *TFRC,* we chose this transcript to test a longer cargo consisting of the subsequent exons 8-19 of *TFRC* (comprising the rest of the CDS) fused to msfGFP, resulting in a 2.1kb cargo (versus the original 791 nt XTEN-GFP cargo). Using this longer TSM, we measured 12% *trans-*splicing efficiency by ddPCR, suggesting that longer cargoes may incur an efficiency trade-off but that *trans-s*plicing of sequences longer than the native CDS is possible (**Fig. S6B**). Taken together, these findings illustrate that our system can install *trans*-spliced cargo into desired locations in diverse endogenous target transcripts with a range of efficiencies and that characteristics of the target intron and cargo modulate *trans*-splicing efficiency.

To investigate the transcriptome-wide specificity of the *trans-*splicing reaction, we selected the top three target transcripts by on-target efficiency (*ITGB1*, *TFRC*, and *SMARCA4*) and used RNA-seq to map all reads containing the TSM cargo. Effective *trans*-splicing was detected for all three targets (36.7%, *ITGB1*; 31.6%, *TFRC*; and 4.2% for *SMARCA4*) (**Table S4**). Importantly, the RNAseq analysis for *ITGB1* confirmed that the combination of TSM with the dCasRx effector and the full CIM with Cas7-11 lead to a significantly higher *trans*-splicing efficiency than either component alone (**Fig. 3F**). The same was true for *trans*-splicing specificity, where we observed a significant 3-fold increase in transcriptome-wide specificity when adding the dCasRx TSM effector compared to the condition with an RNA-only TSM and a Cas7-11 CIM (**Fig. 3F, Fig. S7**).

Overall, we detected a relatively high fraction of total *trans*-splicing occuring on non-target transcripts. When analyzing the distribution of *trans*-splicing across targets however, we observed that for any given off-target transcript, the amount of off-target *trans*-splicing was very low compared to the *ITGB1* target (182 normalized on-target reads; 36.6% on-target efficiency) **(Fig. 3F, 3G, Fig. S7, Table S4**). 77.5% of all detected off-targets across both replicates had an efficiency of less than 5%, indicating that over 95% of the off-target transcripts remained unaffected (**Table S4**). Of the remaining 22.5% of the off-target sites, 86.7% were expressed at extremely low levels (fewer than 10 *cis* and *trans* total normalized junction reads), which would logically result in an outsized contribution of even a single *trans*-spliced read to the observed *trans:cis*+*trans* ratio (**Table S4**). An average of 91.4% of all off-target sites observed in the best *ITGB1* condition were represented by fewer than 10 normalized *trans*-spliced reads, which would indicate that the resultant *trans-*spliced transcript is present at lower abundance than 66.5% of all splice junctions in the transcriptome (**Fig. 3G, Table S4**, **Methods**). Additionally, the overlap of off-target transcripts between replicates was surprisingly low: 22.3% in the presence of TSM guided by dCasRx and Cas7-11 CIM, and 16% in the presence of the TSM guided by dCasRx only. These results reflect the known promiscuity of the spliceosome and indicate a broad off-target profile that is largely stochastic – leading to only subtle overall changes to the cellular transcriptome.

Altogether, these results indicate that the combination of CRISPR-guided transcript targeting and *cis*-splicing interference, and thus the complete RESPLICE system, is critical for efficient and specific *trans*-splicing into endogenous human transcripts.

### RESPLICE replaces pathogenic exons in endogenous human transcripts

Efficient and specific *trans*-splicing could provide a new means to therapeutically target disease-causing transcripts. The technology could correct SNPs, frame-shifts, or repeat expansions by replacing the affected portion of the transcript using a *trans*-splicing reaction directed upstream of the variant location. As an initial proof-of-concept for such future therapeutic applications, here we applied our tool to three endogenous disease-associated transcripts with diverse pathological mechanisms that may benefit from *trans*-splicing manipulations.

Repeat expansions in the first intron of *C9orf72* are responsible for 30% of familial and 5-10% of sporadic cases of Amyotrophic Lateral Sclerosis (ALS) and Frontotemporal Dementia (FTD) (Gijselinck et al., 2018). The nature of the hexanucleotide repeat poses a significant challenge for any gene editing therapeutics (Depienne & Mandel, 2021), and WT C9orf72 protein expression must be maintained to avoid other pathogenic effects (Balendra & Isaacs, 2018). To assess whether *trans-*splicing into the relevant exon of *C9orf72* transcripts with a user-defined cargo was possible, we targeted a TSM encoding a reporter GFP cargo within the first intron (+127), and paired it with a CIM targeting within the same intron (+1152) (**Fig. 4A**). While a targeting TSM with a non-targeting CIM only achieved ∼2% *trans*-splicing efficiency, supplying a targeting CIM achieved ∼10% *trans*-splicing (**Fig. 4A**).

**Figure 4:**
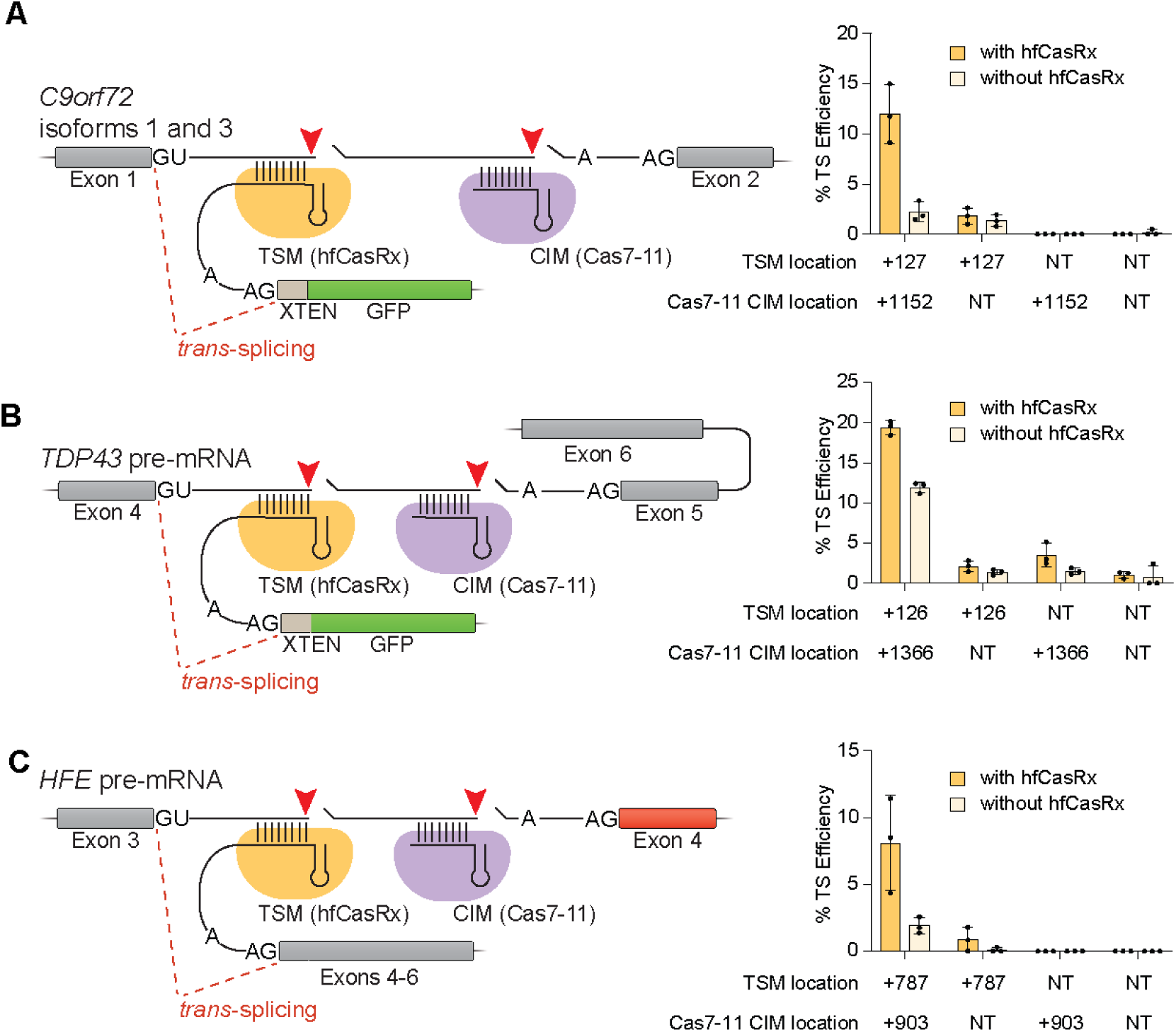
RESPLICE replaces and corrects repeat expansions and deleterious mutations in pathogenic endogenous human transcripts. **(a)** Schematic depicts the desired *trans*-splicing reaction downstream of exon 1 of *C9orf72*, with the TSM, guided by hfCasRx (yellow) , and the Cas7-11 CIM (purple). On the right is a plot showing *trans*-splicing efficiency measured by ddPCR onto exon 1 of *C9orf72* with either the highest efficiency TSM or non-targeting TSM, combined with either the highest efficiency CIM or a non-targeting CIM in HEK293T cells. **(b)** Schematic showing *trans*-splicing downstream of exon 4 of *TDP43*, with the hfCasRx TSM (yellow) targeting intron 4, which is upstream of exons 5 and 6 of *TDP43* that usually contain mutations in the case of disease. The Cas7-11 CIM (purple) targets downstream of the TSM binding site within intron 4. On the right is a plot showing *trans*-splicing efficiency onto exon 4 of *TDP43* with either the highest efficiency TSM or non-targeting TSM, combined with either the highest efficiency CIM or non-targeting CIM in HEK293T cells. **(c)** Schematic of the desired *trans*-splicing reaction downstream of exon 3 of *HFE,* which carries a pathogenic mutation in exon 4 in Huh7 cells. The hfCasRx TSM (yellow) and downstream Cas7-11 CIM (purple) are shown targeting intron 3. In this TSM, the cargo consists of exons 4 through 6 of the HFE transcript to replace the remaining downstream exons with wild-type sequences, aiming to correct the pathogenic mutation to yield wild-type HFE protein (endogenous exons in red). On the right is a plot showing *trans*-splicing efficiency, as measured by ddPCR, downstream of *HFE* exon 3 using either the highest efficiency TSM or non-targeting TSM, in combination with either the highest efficiency CIM or the non-targeting CIM in Huh7 cells. All data are mean ± s.d.; n = 3 biological replicates, with individual points plotted.

*TDP43* encodes a protein implicated in neurodegenerative disorders with a broad spectrum of mutations, largely localized in its glycine-rich domain (Buratti, 2015). The diversity of pathogenic variants makes *TDP43* a difficult target for gene editing and other therapeutic strategies, similar to other disease-associated genes with high mutational heterogeneity such as *RPE65* in the hereditary blindness disorder Leber congenital amaurosis (Hanany et al., 2020), or *CFTR* in cystic fibrosis (Drumm et al., 2012). Here, we designed a *trans*-splicing system to install a reporter cargo downstream of *TDP43* exon 4, upstream of the glycine-rich domain – aiming to simulate a one-size-fits-most approach for restoring full-length functional protein expression by replacing mutated downstream exons in a variant-agnostic manner (**Fig. 4B**). We observed 20% *trans*-splicing efficiency of a reporter cargo at the intended site when using targeting versions of both the TSM (+126) and CIM (+1366) (**Fig. 4B**). We then replaced the msfGFP cargo (791 nt) with the remaining coding sequence of *TDP43* (exons 5 and 6, 702 nt) to closely mimic a therapeutic cargo and demonstrated 3.8% *trans*-splicing efficiency (**Fig. S6B**).

Hereditary hemochromatosis – an autosomal recessive genetic disorder resulting in deposits of excess iron that cause multi-organ dysfunction – is caused by multiple point mutations in the gene *HFE*, most of which occur downstream of exon 3 (**Fig. 4C**) (Wallace & Subramaniam, 2016). The only current treatment option for primary hemochromatosis patients is lifelong, frequent phlebotomy to deplete red blood cells and alleviate iron toxicity (Bacon et al., 2011). The Huh7 cell line natively contains an in-frame deletion of *HFE* Tyr271, which has been shown to prevent HFE membrane localization and compromise its function (Vecchi et al., 2010). We replaced the synthetic cargo from our other endogenous targets with a 457 nt therapeutically-relevant cargo encoding *HFE* exons 4 and 5, and demonstrated 8% *trans*-splicing efficiency. Importantly, this result demonstrates the utility of our *trans*-splicing system in a second human cell line. While optimization of the system is likely to improve efficiency further, it is notable that an *in vivo* CRISPR base editing preclinical study in a mouse model of hereditary hemochromatosis demonstrated significant alleviation of iron toxicity and therapeutic benefit with 10.7% base editing-mediated correction of *HFE* C282Y (Rovai et al., 2022). Therefore, our *trans*-splicing efficiency of 8% would also be expected to yield detectable therapeutic benefit in a comparable model system, while also enabling the treatment of a broad set of pathogenic variants and avoiding concerns of permanent genomic alterations that can arise from DNA-targeted gene editing systems.

Altogether, we show that our *trans-*splicing system can be readily applied to disease-associated endogenous transcripts that represent several classes of pathogenic mutations uniquely well-suited to splicing-mediated intervention – and currently lacking robust therapeutic options. Our transcriptome-wide analysis of *trans*-splicing into reporter and endogenous human transcripts indicates a strong potential for our programmable system, RESPLICE, to generate efficient and specific large-scale edits to target transcripts of interest, substantially broadening the RNA editing toolbox.

## DISCUSSION

In this study, we report our development of a programmable *trans*-splicing system, RESPLICE, that can efficiently splice exogenous cargo exon(s) onto a CRISPR-dCasRx-specified splice donor site in endogenous transcripts – a first for the field of *trans*-splicing. RESPLICE uses two RNA-targeting modules: the TSM to localize the *trans*-splicing reaction, and the CIM to inhibit the endogenous *cis*-splicing reaction. Strikingly, we were able to achieve up to 47% *trans*-splicing of a fluorescent cargo onto endogenous transcripts in unsorted HEK293T cells with high specificity, as dictated by the programmable CRISPR effectors. Together, we demonstrate efficient, specific, and programmable *trans*-splicing in two human cell types across nine endogenous transcripts, which paves the way toward exciting basic science applications (e.g. transient labeling or relocalization of endogenous proteins) and therapeutics (e.g. targeting repeat expansion disorders, aberrant splicing patterns, or multiple pathogenic variants in the same protein).

Our study demonstrates how the combination of our TSM and CIM modules enables efficient *trans*-splicing into endogenous transcripts. Previous work in the *trans*-splicing literature primarily relied on fluorescent reporters, gel electrophoresis of RT-PCR products, or western blots of protein products to assess efficiency and specificity (Kikumori et al., 2001; Mansfield et al., 2000; Puttaraju et al., 1999; Tockner et al., 2016). For this reason, it is challenging to directly compare our efficiencies to these previous studies. As we were preparing this manuscript, a preprint reporting a CRISPR-Cas13-mediated *trans-*splicing system achieved up to 60% of gated cells with detectable *trans*-splicing by flow cytometry in a fluorescent mini-gene reporter system (Borrajo et al., 2023). While conceptually similar to our TSM module, this system did not apply any means to disfavor the *cis-*splicing reaction, which we show to be critically important for efficiency when targeting endogenous human transcripts – a target type that was not explored in this preprint. Despite these and other key differences, both of our works demonstrate that RNA-targeting CRISPR effectors can enhance the efficiency of *trans*-splicing, and open up many new opportunities for further technology development in this space.

The spliceosome is inherently a promiscuous RNA editing machine, relying on the integration of signals from many *cis*- and *trans*-acting factors to define exons and introns of each pre-mRNA (Matera & Wang, 2014). Therefore, splicing is a noisy, variable process resulting in many erroneously spliced transcripts which are degraded by multiple RNA surveillance mechanisms (Wolin & Maquat, 2019). Indeed, our deep, transcriptome-wide assessment of *trans*-splicing events – the only such profiling that has been reported in any *trans*-splicing manuscript, to our knowledge – supports this notion: we observed very low-level off-target *trans*-splicing into hundreds or thousands of different splice sites, often barely detectable at fewer than 10 normalized reads. We do not believe that this level of off-target *trans*-splicing would functionally impact the cell, as these changes are transient and impact a small percentage of the copies of each transcript in question. Analyses of splicing fidelity have found that errors in splicing occur at a rate greater than 0.1% per intron on average (Pickrell et al., 2010; Skandalis, 2016) which is substantially higher than the rate of *trans*-splicing off-targets we measured, occuring at less than 0.005% on average of all splice junctions across all of our RNA-seq samples (**Table S5**). This suggests that off-target *trans*-splicing happens much less frequently than the basal splicing error rate and should therefore have minimal effects on the cell. Most, if not all, genome editing systems are limited to some extent by off-targets, with DNA cleavage off-targets being the most deleterious.

When targeting endogenous human transcripts, we found that *trans-*splicing specificity and efficiency improved drastically with the inclusion of dCasRx to guide the TSM to improve specificity and a catalytically-active Cas7-11 CIM to interfere with the *cis*-splicing reaction. We suspect that cleavage of the target transcript within the intron may stimulate its ability to be *trans*-spliced. Furthermore, our data across 9 endogenous human transcripts indicated that target introns between 0.7-6kb seem to be optimal for higher *trans-*splicing efficiency. Additionally, our evidence anecdotally suggests that features of the cargo, such as length and sequence, can have a substantial effect on *trans*-splicing efficiency, as seen when swapping TSM cargoes for *TFRC* and *TDP43* (**Fig. S6**). Overall, we hypothesize that many factors such as intron strength, intron length, gene structure, topological location, and/or surrounding sequence may have a significant effect on efficiency and specificity. Future work comprehensively testing RESPLICE across many endogenous transcripts will be necessary to identify the key design features contributing to highly efficient and specific *trans*-splicing.

For research applications, *trans*-splicing could enable significantly easier functional characterization of splice isoforms. We showed in the RNA-seq analysis of *SMARCA4* that RESPLICE can successfully edit endogenous alternatively spliced transcripts. A splice isoform of interest can be expressed in a tunable manner by targeting the upstream splice donor and supplying a TSM with the corresponding cargo. Beyond splice isoform characterization, RESPLICE can install desired protein fusions (ie. msfGFP, FLAG tag, etc.) or sequences (ie. different 3’ UTRs, localization sequences, fluorescent aptamers, different poly-A signals, etc.) to study and modify a transcript of interest without significantly perturbing its expression level. Given that this modification occurs at the RNA level, it also has the potential to be temporally controlled and reversible.

For therapeutic applications, *trans*-splicing could be particularly useful as an intervention in diseases caused by genetic rearrangements, splice aberrations, or repeat expansions. It provides a means to restore functionality to particularly long genes that are difficult to introduce by traditional gene replacement therapy (e.g. dystrophin). Moreover, *trans*-splicing could enable one-size-fits-most therapies when mutational heterogeneity in the pathogenic gene would otherwise necessitate dozens of site-specific gene editing approaches for different patients (e.g. Leber congenital amaurosis, cystic fibrosis), or in repeat expansion disorders (e.g. spinocerebellar ataxia, ALS). The simultaneous knockdown and replacement at a therapeutically relevant junction provides a proof-of-concept for future development of *trans*-splicing-based therapeutics to replace expanded repeats in *C9orf72* and possibly other repeat expansion disorders.

Finally, the transient nature of RNA-based therapies and RNA editing alleviates many of the safety concerns that come with DNA-targeted gene editing therapeutics that are entering the clinic. In this new era of RNA medicine, our work now provides the first opportunity to make large-scale changes at robust efficiencies to the sequence of endogenous human transcripts, providing a foundation for future biotechnological and therapeutic development of *trans*-splicing technology.

**Table S1**

**Table S2**

**Table S3**

**Table S4**

**Table S5**

## METHODS

### Design and cloning of bacterial constructs

The reporter plasmid consists of the 5’ half of the msfGFP coding sequence, followed by intron 4 of chicken cardiac troponin and an artificial exon from the RG6 minigene in a standardized plasmid expression backbone, driven by an EF1a promoter (Konermann et al., 2018; Orengo et al., 2006).

For our reporter assay, the TSM plasmid contains a CasRx or Cas7-11 direct repeat (DR), a type IIS landing pad site, a stuffer sequence (Uckun et al., 2015), splice motifs (Uckun et al., 2015), the 3’ half of the msfGFP coding sequence, a T2A ribosomal skip sequence, and full-length mTagBFP2 in Puc19 (Addgene Plasmid #50005), driven by a SFFV promoter (from plasmid #pFu-62 derived from Addgene Plasmid #57827, gift from Becky Xu Hua Fu). TSM spacers were sub-cloned into this plasmid using Golden Gate cloning. For our promoter screening, we used Gibson Assembly to subclone the above plasmid to contain the EF1a, EFS, or CMV promoters.

The TSM effector plasmids contained human codon-optimized sequences for catalytically inactive dCasRx (Konermann et al., 2018), high-fidelity catalytically active hfCasRx (Tong et al., 2022), or catalytically active Cas7-11 (Özcan et al., 2021) followed by the P2A ribosomal skip sequence, full-length mCherry, and a WPRE sequence in pLentiGuide-Puro (Addgene Plasmid #52963), driven by the EF1a promoter. For TSM effector plasmids containing the dsRBD fusion, a sequence encoding the human codon-optimized first 80 amino acids of first double-stranded RNA binding motif of eukaryotic translation initiation factor 2-alpha kinase 2 were added to the C-terminal end of dCasRx or hfCasRx, before the P2A sequence in accordance with the best construct designed previously (Han et al., 2020).

For our endogenous transcript *trans*-splicing experiments, the same TSM plasmid as above was subcloned to change the cargo to the XTEN16 linker followed by full-length msfGFP, or human codon-optimized amino acids 206-348 of *HFE,* or human codon-optimized amino acids 182-414 of TDP43, or human codon-optimized amino acids of TFRC followed by the XTEN16 linker and full-length msfGFP using Gibson Assembly. TSM spacers were subcloned using Golden Gate Assembly, as described above.

For our CIM all-in-one plasmid, we subcloned the TSM effector plasmids above to also contain a U6 promoter with the corresponding DR (CasRx or Cas7-11) and a Type IIS landing-pad using Gibson Assembly. For our reporter assay, we generated CIM all-in-one plasmids for hfCasRx, hfCasRx-RBD, and Cas7-11. For endogenous experiments, we subcloned the Cas7-11 all-in-one plasmid to replace mCherry with mTagBFP2. We sub-cloned the CIM spacers using Golden Gate Assembly.

All molecular cloning steps were transformed into Mach1 cells.

### Mammalian cell culture and transient transfection

HEK293T (ATCC# CRL-3216) and Huh-7 cells were obtained from the Berkeley Cell Culture facility. HEK293T cells were cultured in a controlled humidified incubator at 37 °C and 5% CO2 in DMEM (Gibco) media supplemented with 10% FBS (Gibco 26140095), penicillin (10,000 IU mL-1) and streptomycin (10,000 ug mL-1). Huh7 cells were cultured in the same incubator conditions (37 °C, 5% CO2) in low glucose DMEM (Gibco) media supplemented with 10% FBS (Gibco 26140095), penicillin (10,000 IU mL-1) and streptomycin (10,000 ug mL-1) and sodium pyruvate (110 mg L-1).

For transient transfections for flow cytometry, reverse transcription, and RNAseq experiments, HEK293T cells were seeded at a density of 16k cells per well and Huh7 cells were seeded at 5k cells per well in 96-well TC treated plates at most 16 hours before transfection. Each condition was performed with 3 biological replicates. For transient transfections for sorting experiments, HEK293T cells were seeded at a density of 150k cells per well in TC treated 6-well plates at most 16 hours prior to transfection. Each condition was performed with 2 biological replicates. All plasmids used in transfections were prepared using the Nucleobond Xtra Midi EF Kit (Machery Nagel) according to the manufacturer’s protocol.

For the reporter assay without CIM, 100 ng of TSM plasmid, 100 ng of reporter plasmid, and 100 ng of TSM effector plasmid per well were transfected with Lipofectamine 2000 following manufacturer’s protocol. For the reporter assay with CIM, 100 ng of TSM plasmid, 100 ng of reporter plasmid, 100 ng of TSM effector plasmid, and 100 ng of CIM plasmid were transfected using Lipofectamine 2000. For endogenous assays without CIM, 200 ng of TSM plasmid and 200 ng of TSM effector plasmid per well were transfected using Lipofectamine 2000. For endogenous transcript targeting assays with CIM in HEK293T cells, 200 ng of TSM plasmid, 100 ng of TSM effector plasmid and 200 ng of CIM plasmid per well were transfected using Lipofectamine 2000. For endogenous transcript targeting assays with CIM in Huh-7 cells, 200 ng of TSM plasmid, 100 ng of TSM effector plasmid and 200 ng of CIM plasmid per well were transfected using Lipofectamine 3000, following the manufacturer’s protocol. For the sorting experiment, 6 ug of TSM plasmid, 3 ug of TSM effector plasmid, and 6 ug of CIM plasmid per well were transfected using Lipofectamine 2000. For all experiments containing no effector controls (for either TSM or CIM), the same mass of pUC19 was added in lieu of the effector plasmid such that the total mass of DNA remained constant.

### Flow cytometry

After 48 hours following transfection, the media was removed from each well of the 96-well plate. 50 µL of TrypLE Express was added to each well and the plate was incubated at 37 °C for 7 minutes. After incubation, 100 µL of stain buffer (Invitrogen) was added and mixed vigorously. This mixture was then transferred to a 96-well U-bottom plate (Falcon) and the plate was then spun at 300 x g for 5 minutes. The supernatant was carefully removed without disturbing the cell pellet, and 200 µL of stain buffer was then added and mixed vigorously. All flow cytometry was performed in 96-well format with the same Attune NxT Flow Cytometer (ThermoFisher) and analyzed using FlowJo. Each sample contains at least 50,000 total events and gates were set using un-transfected and single color controls (**Fig. S8**).

### Lysis, RNA extraction, and reverse Transcription for qPCR and ddPCR assays

After 48 hours following transfection, cells were lysed and reverse transcribed according to a previously developed protocol (Joung et al., 2017). 3 µL of RNA lysis stop solution (1 mM Proteinase K inhibitor, 90 mM EGTA, and 113 μM DTT in UltraPure water) was aliquoted into each well of a 96-well PCR plate. After removing cell culture media and washing with DPBS (without Ca or Mg) (Gibco), 50 µL of RNA Lysis Buffer (9.6 mM Tris–HCl (pH 7.8), 3 U ml−1 Proteinase K, 300 U mL-1 DNAse 1, 0.5 mM MgCl2, 0.44 mM CaCl2, 10 μM DTT, 0.1% (wt/vol) Triton X-114, and 3 U ml−1 proteinase K) were added to each well of the 96-well tissue culture plate and incubated at RT for 10 minutes. Cells and RNA lysis buffer were mixed and 30 µL of the lysed cells were transferred into the 96-well PCR plate with aliquoted stop lysis buffer and mixed and incubated for 3 minutes to stop the lysis reaction. 32 µL of RevertAid RT Reverse Transcription Kit using random hexamer primers (Thermofisher) prepared using the manufacturer’s protocol was aliquoted into a separate 96-well PCR plate and 8 µL of the lysis samples were added and mixed. The reverse transcription mix was incubated at 25 °C for 10 minutes, 37 °C for 60 minutes, and 95 °C for 5 minutes and either used immediately or frozen at -80 °C.

### Quantitative and digital droplet PCR and probe primer design

We designed all *cis*- and *trans*- splicing qPCR and ddPCR primers using Primer-BLAST to minimize off-target hybridization to the transcriptome (Ye et al., 2012). All PCR products were designed to have an amplicon length of 70-200 bp and melting temperatures between 57°C and 63°C and contain at least 3 mismatches to any off-targets. Probes were designed on the *cis*- or *trans*- splicing junction with melting temperatures under 37°C on either side of the junction and with an overall melting temperature of > 55°C.

### Quantitative PCR

A master mix was made using TaqMan Fast Advanced qPCR mix and 3 different primer-hydrolysis probe mixes. One primer-probe mix was designed to detect the expected corresponding *trans*-splicing junction of the target gene, the next used the pre-designed PrimeTime qPCR Probe Assay to detect the exons 4-5 boundary of GAPDH, and the final mix used previously designed primers and probes to detect an upstream exon junction of the target gene. 15 µL of the qPCR master mix was added to 9.6 µL of the reverse-transcription reaction from above, and qPCR was carried out in 5 µL multiplexed reactions and 384-well format using the LightCycler 480 Instrument II (Roche). Final concentration of primers are 0.5 µM and probes are 0.25 µM. First, we filtered out replicates where the delta Cp between GAPDH and either the *trans*-splicing junction or upstream exon was quite different from the other replicates. Next, the delta Cp between the junction and upstream exon was calculated and then averaged across all filtered replicates corresponding to the same biological replicate. The junction and upstream exon delta cp was converted into a % on-target *trans*-splicing efficiency by raising 2 to the power of the delta Cp and multiplying by 100.

### Digital droplet PCR

A master mix was made using ‘ddPCR Supermix for Probes (no dUTP)’ and 2 different primer-probe pairs. One primer-probe mix used sequences from PrimeTime qPCR Probe Assays from IDT for GAPDH or TBP, while the other was designed to either target the trans-splicing junction or cis-splicing junction of the target. The final concentration of primers and probes were 0.9 µM and 0.25 µM respectively. For all target genes except for C9ORF72, HFE, and TDP43, the reverse transcription reaction was diluted 1:20 in water, and 1.1 µL of that dilution was added to 20.9 µL of the ddPCR mastermix with GAPDH as the housekeeping gene. For TDP43, the reverse transcription reaction was diluted 1:5, and 1.1 µL of that dilution was added to 20.9 of the ddPCR mastermix with TBP as the housekeeping gene. For C9ORF72 and HFE, the reverse transcription reaction volume was doubled, and the resulting cDNA was extracted using bead cleanup with a 1.8:1 ratio of beads to reverse transcription reaction, according to the manufacturer’s protocol. Following two washes with 80% ethanol, the cDNA was eluted in 12 µL of water. 5 µL of this extraction was added to 17 µL of the mastermix with TBP as the housekeeping gene, where the final TBP primer concentrations were 0.45 µM and probe concentration was 0.125 µM. The assembled ddPCR reactions were then loaded into the Automated Droplet Generator. Following droplet generation, the ddPCR reactions were cycled in thermal cycler for 1 cycle for 30 minutes at 90C, 40 cycles with an annealing temperature of 54C for 1 minute followed by an extension at 72C for 1 minute, and 1 cycle for 10 minutes at 90C, followed by 4C infinity step. After amplification, the ddPCR reactions were sealed and read using the Biorad Digital Droplet Reader, which calculates the copies/µL of the probed transcripts from the number of positive droplets per well. The concentration of cis or trans-junctions were normalized to the housekeeping gene, and the percent trans-splicing efficiency was calculated by dividing the normalized trans-junction concentration by the normalized sum of cis and trans-junction concentration and multiplied by 100. The relative expression of each gene was calculated by dividing the normalized total concentration by either the targeting TSM and non-targeting CIM control for the ITGB1 CIM tiling experiment, or the non-targeting TSM and non-targeting CIM control for all other experiments.

### Sorting

After 48 hours, the media was removed from each well of the 6-well plate. 1 mL of TrypLE Express was added to each well and the plate was incubated at 37°C for 7 minutes. After incubation, 1 mL of stain buffer (ThermoFisher Sci 00-4222-57) was added and pipette mixed thoroughly. The cells were transferred into 2 mL microcentrifuge tubes and spun down at 200g for 5 minutes. The supernatant was carefully removed without disturbing the cell pellet, and 1 mL of stain buffer was added and mixed thoroughly to resuspend the pellet. The cells were strained through a 5mL Falcon tube cell strainer snap cap prior to sorting (Corning #352235). All sorting experiments were carried out on the BD FACSAria Fusion. Live and single cell gates were set on untransfected cells using forward and side scatter, and dCasRx effector positive cells were selected for by gating for mCherry positive cells. These cells were then binned into 4 quadrant bins by BFP expression (**Figure 3D(i)**). ∼300,000-500,000 cells were collected for each bin across each condition in 5 mL collection tubes filled with 500 µL of cold stain buffer. Following sorting, the cells were transferred from the 5 mL collection tubes to 1.5 mL microcentrifuge tubes and spun down at 100g for 10 minutes. Once the supernatant was carefully removed and the pellet was visually confirmed, 100 µL of lysis buffer was added to each cell pellet and mixed vigorously. Following a 10 minute incubation, 60 µL of the lysis was transferred into a PCR-plate containing 6 µL of stop-lysis solution, and incubated for 3 minutes. Then, the samples were reverse transcribed and quantified using ddPCR as previously described.

### RNA extraction, library preparation, and RNA-Seq

After 48 hours, two replicates of each condition were extracted using the NucleoSpin RNA Plus kit (Machery Nagle) and used on-column DNAse digestion following the manufacturer’s protocol. 750 ng of total RNA was used for each sample to prepare libraries with the NEBNext Ultra II Kit with 8 cycles of amplification. The samples were sequenced using the Illumina NovaSeq-X with 150 bp paired end reads.

### RNA-seq analysis

Sequenced reads were trimmed for adapters and filtered by length (≥ 40 bp) and quality (Q-score ≥ 10) using CutAdapt v4.4. Next, reads were aligned using STAR aligner v2.7.3a (Dobin et al., 2013) to a custom index containing hg38 human genome and a sequence corresponding to the target upstream exon, the target intron and the *trans*-spliced cargo. For samples with the reporter, the index consisted of the hg38 genome, one sequence containing the first exon of the reporter followed by the intron and reporter second exon, and a second sequence containing the reporter first exon, intron, and reporter *trans*-spliced cargo. Mapping was carried out using a max mismatch of 4 bp for all parameters, a minimum chimeric segment and splice junction overhang of 20 bp, and a maximum intron length of 100,000 bp. On average ∼145M reads were mapped for each sample.

*Cis*-spliced and on-target *trans*-spliced junctions were counted in the splice-junction output file generated by STAR for each sample. Off-target *trans*-splicing junctions were found in the Chimeric Splice Junction Alignment file output from STAR. The first filtering step isolated reads mapping to the first nucleotide of the *trans*-splicing cargo. Next, the reads were trimmed to remove any part mapping to the *trans*-splicing cargo. The trimmed reads were then mapped to the hg38 human genome using standalone Nucleotide-Nucleotide BLAST v2.14.1+, and any reads that did not map to the human genome were filtered out. The highest quality map was then defined as the off-target *trans*-splicing site and the number of reads corresponding to *cis*- and *trans*-splicing of the off-target splice junction were recorded. With these counts, we normalized each to the average number of reads mapped across samples (∼145M). With these normalized counts, we only included *trans*-splicing events where the corresponding *cis*- or *trans*-spliced junctions had counts ≥ 3.0 normalized reads, a previously established cutoff (Emig et al., 2010).

On-target % *trans*-splicing efficiency was calculated by taking the ratio of filtered and normalized on-target *trans*-splicing junction reads to the sum of normalized and filtered *cis*- and *trans*-spliced junction reads that share the same target splice donor and multiplying by 100. On-target % *trans*-splicing specificity was calculated by taking the ratio of filtered and normalized on-target trans-splicing junction reads to the total number of filtered and normalized on- and off-target *trans*-splicing reads. The number of total off-target *trans*-splicing junctions was calculated by counting the number of splice junctions present in all samples within a comparison (each junction only counted once). The number of overlapping off-target *trans*-splicing junctions was calculated by counting the number of splice junctions that were present in each sample within a comparison. Off-target *trans*-splicing genes were found by mapping the coordinate of the off-target *trans*-splicing splice donor to a gene using the PyEnsembl package with the Ensemble 110 Release. The number of total and overlapping off-target *trans*-splicing genes was calculated the same as splice-junctions above. We analyzed the distribution of the number of splice junctions pUC-only transfected HEK293T controls and found that an average of 66.5% of the filtered splice junctions had normalized junction read counts greater than 10, indicating junctions with normalized counts below 10 reads are low abundance.

For the reporter samples, two alternative splice donors were identified in the bichromatic splice reporter in addition to the intended splice donor. Each of these splice donors were capable of splicing with the TSM, so to maintain consistency with the flow cytometry readout, which can only identify splicing into the intended splice donor with reconstituted GFP, and since the other isoforms utilized different splice donors, we only included the reconstituted GFP splice junctions when calculating efficiency and specificity (**Fig. S3**). For *SMARCA4*, a second isoform was found in the RNA-Seq analysis, and its counts were included in the calculation of on-target *trans*-splicing efficiency since it shared the sample splice donor as the intended isoform (**Table S4**).

### Statistical analysis

All values are reported as mean ± SD as indicated in the appropriate figure legends. For correlation analysis, Pearson’s correlation coefficient was used. Two-tailed Welch’s test was used for statistical comparisons. The Benjamini-Hochberg multiple testing p-value correction procedure was used for the statistical analysis of the reporter CIM tiling screen. Statistical functions from the scipy package (v1.11.4) were used for all statistical analyses. Three biological replicates were used for every experiment besides RNA-seq where only two replicates were used.

### Data and Software Availability

Data and software used in this study are available at https://github.com/hsulab-arc/RESPLICE

## Acknowledgements

We thank the entire Hsu lab for helpful discussions and feedback. We also thank the Arc Institute Scientific Publications Team for assistance with the manuscript, the Arc Multiomics Center and Genomics Platform for assistance with RNA-seq, B. Xu Hua Fu for providing a plasmid and helpful discussions, A. Sclip for feedback on the manuscript and helpful discussions, and J. McSpedon for advice and assistance with RNA-seq analysis. S.S.C. is supported by the Paul and Daisy Soros Fellowship, the Siebel Scholarship and the Arc Institute. C.T., M.N., and S.K. are supported by funding from the Arc Institute. P.D.H. is supported by funding from the Arc Institute, Rainwater Foundation, Curci Foundation, Rose Hill Innovators Program, V. and N. Khosla, and anonymous gifts to the Hsu Lab.

## Author Contributions

S.S.C., C.T., and P.D.H. conceived the study. S.S.C., C.T., and M.N. performed experiments. C.T. performed computational analyses. S.S.C., C.T., S.K., and P.D.H. designed the experiments and analyzed the results. P.D.H. and S.K. supervised the research. S.S.C., C.T., A.P, S.K., and P.D.H. wrote the manuscript.

---

**Supplementary Figure 1.**
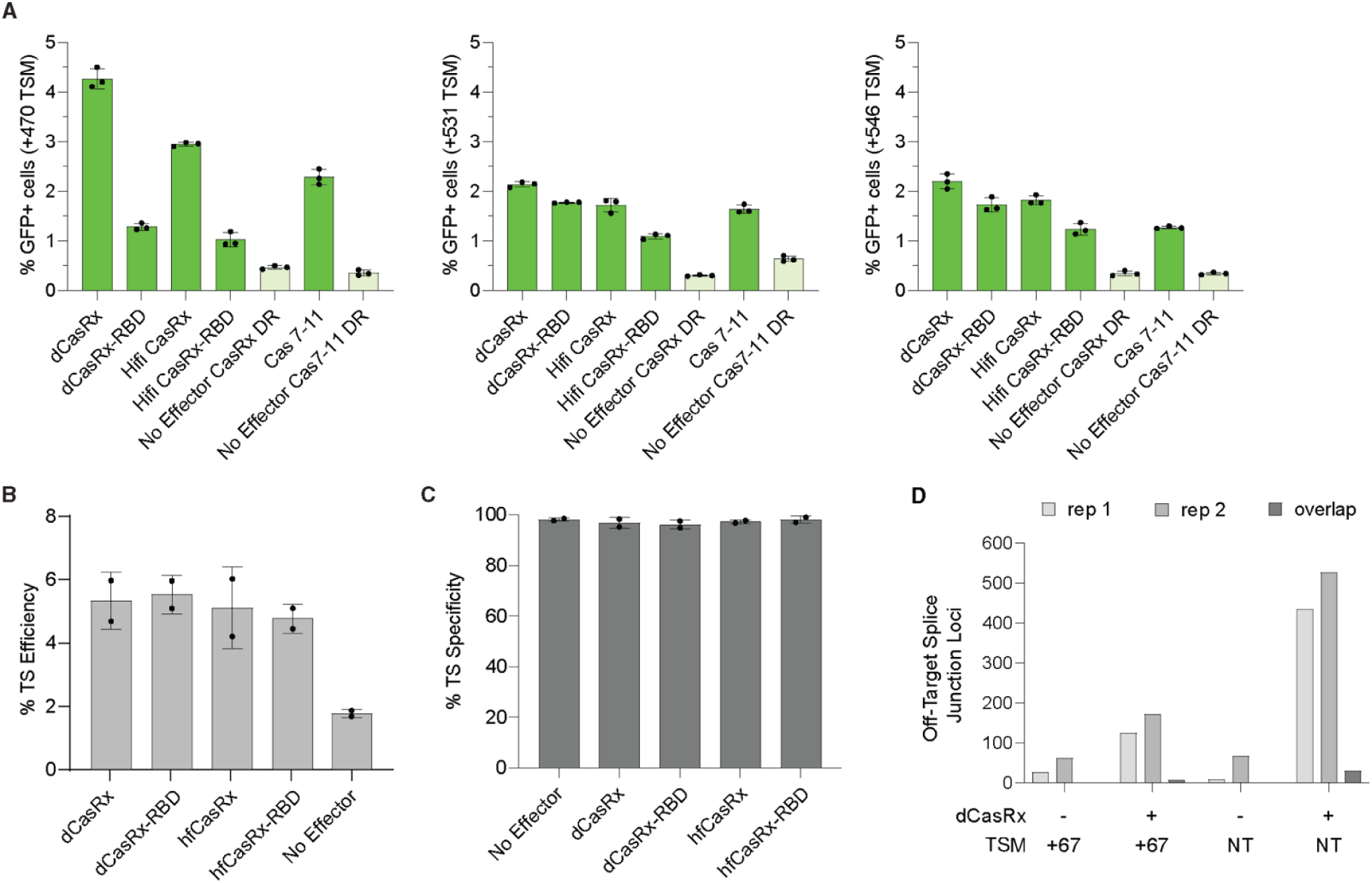
Comparison of TSM Effectors by Flow Cytometry and RNAseq. **(a)** A comparison of efficiency and specificity across different effectors and TSM targeting locations. Percentage of single HEK293T GFP^+^ cells are plotted on the left y-axis in green bars for the given TSM spacer (470 nucleotides downstream of the splice donor for the leftmost plot, 531 nucleotides downstream of the splice donor for the middle plot, 546 nucleotides downstream of the splice donor for the rightmost plot). Data are mean ± s.d.; n = 3 biological replicates. **(b)** A comparison of the *trans*-splicing efficiency of different effectors using RNA-seq in HEK293T cells. Data are mean ± s.d.; n = 2 biological replicates, with individual points plotted. **(c)** A comparison of the *trans*-splicing specificity of different effectors using RNA-seq in HEK293T cells. Data are mean ± s.d.; n = 2 biological replicates, with individual points plotted. **(d)** A comparison of the number of individual and overlapping off-target *trans*-splicing junction loci with and without dCasRx for both targeting and NT TSMs in HEK293T cells.

**Supplementary Figure 2.**
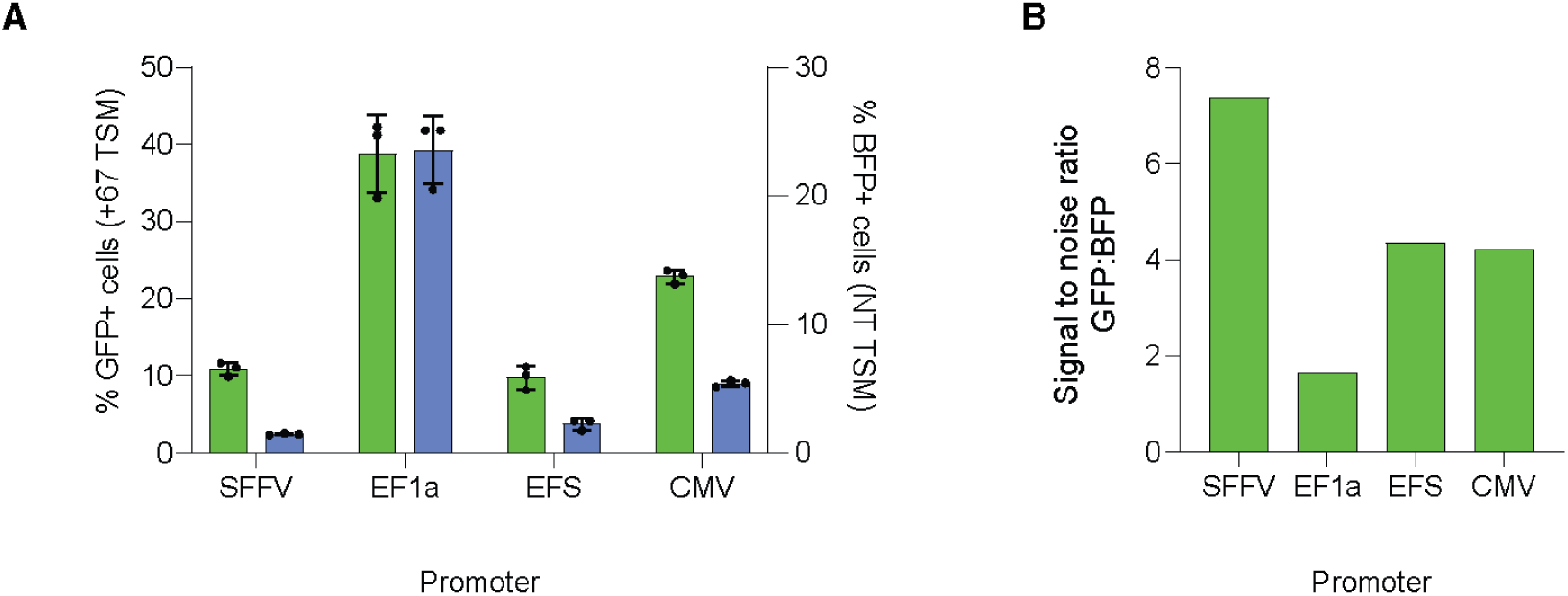
Comparison of Promoters for TSM Expression. **(a)** A comparison of efficiency and specificity across different promoters driving the TSM. Percentage of single HEK293T GFP^+^ cells are plotted on the left y-axis in green bars for the TSM spacer targeting 97 nucleotides downstream of the splice donor on the reporter transcript. Percentage of single HEK293T BFP^+^ cells are plotted on the right y-axis in blue bars for the non-targeting TSM spacer. Data are mean ± s.d.; n = 3 biological replicates, with individual points plotted. **(b)** A comparison of the signal to noise ratio for each promoter which is calculated by dividing the mean percentage of single HEK293T GFP^+^ cells with the +97 TSM by the mean percentage of single HEK293T BFP^+^ cells with the non-targeting TSM across three biological replicates.

**Supplementary Figure 3.**
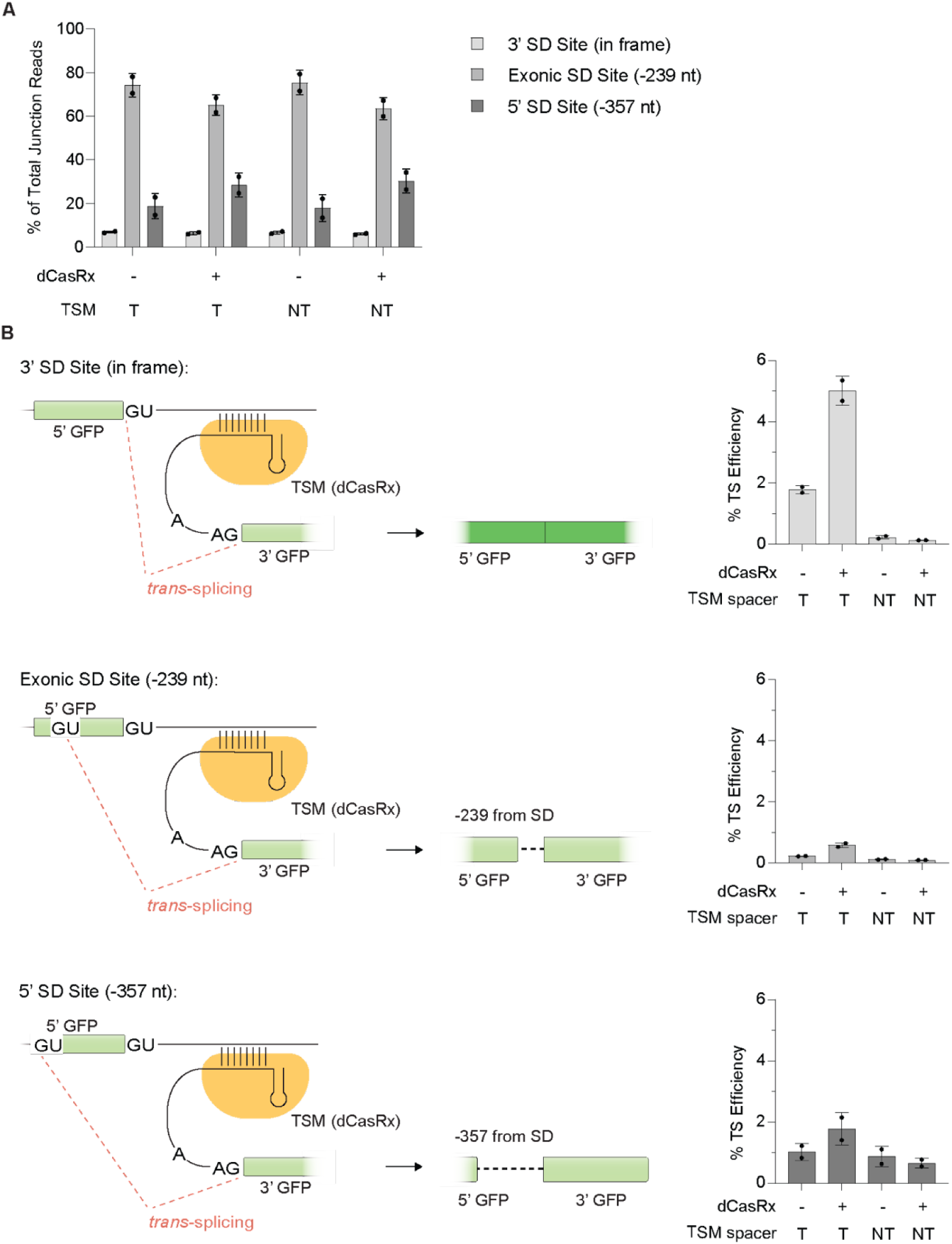
Assessment of Alternative Splicing of Bichromatic Reporter. **(a)** Analysis of splice isoforms for the reporter transcript using RNA-Seq in HEK293T cells. Data are mean ± s.d.; n = 2 biological replicates, with individual points plotted. Total junction reads are the sum of the *cis* and *trans* junction reads at each junction site. **(b)** Schematics and the associated *trans*-splicing efficiencies for the different splice isoforms of the reporter in HEK293T cells. Data are mean ± s.d.; n = 2 biological replicates, with individual points plotted. *Trans*-splicing efficiency is calculated as percentage of *trans* junction reads out of total junction reads (sum of *cis* and *trans* junction reads).

**Supplementary Figure 4.**
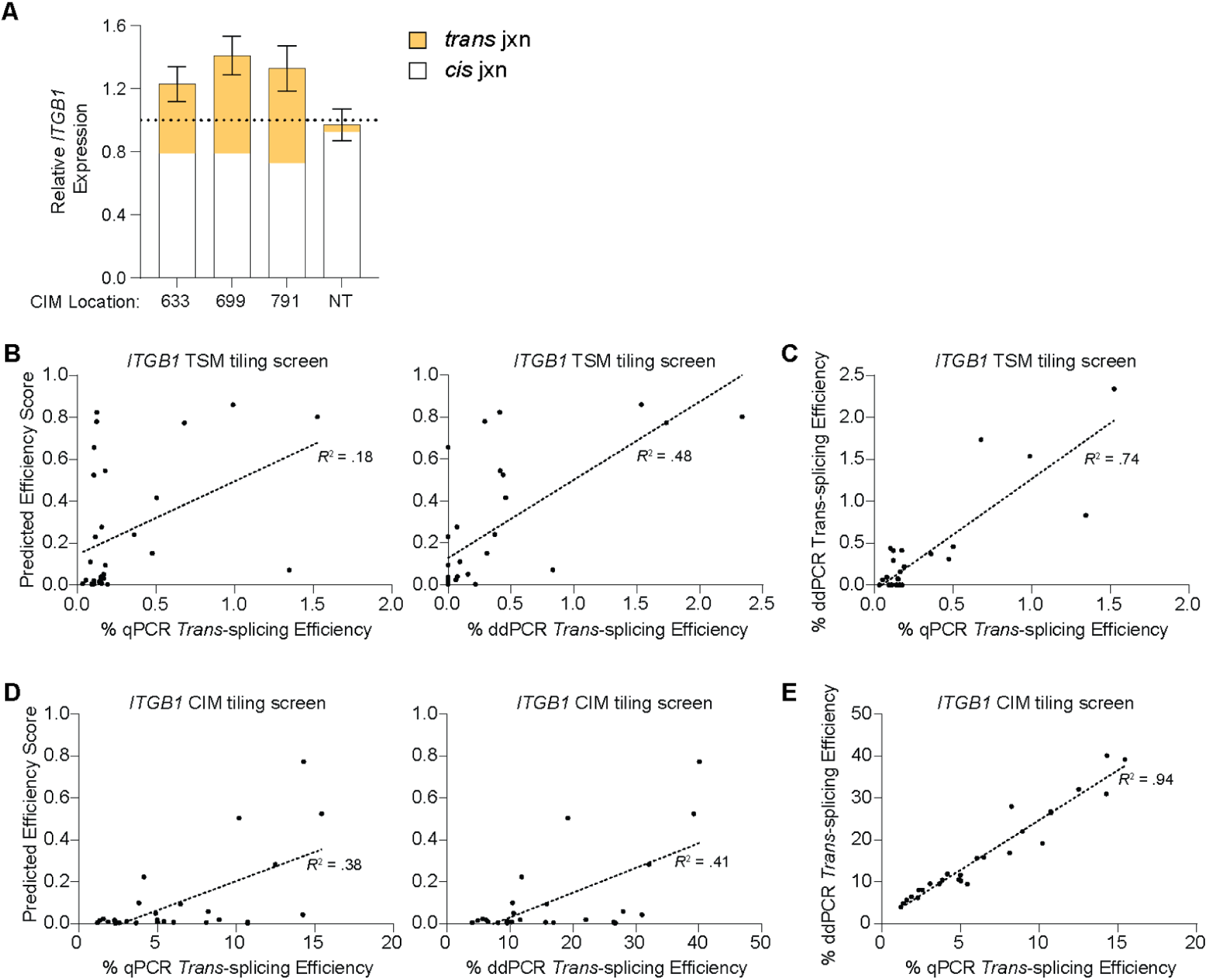
Analysis of TSM and CIM Spacer Efficiencies. **(a)** Fold change of *ITGB1* expression averaged across three replicates of each Cas7-11 CIM (+633 nt, +699 nt, +791 nt, NT relative to the NT CIM for the +231 TSM spacer measured by ddPCR. The gray portion of the bar corresponds to the proportion of *cis* junctions, while the yellow portion of the bars correspond to the proportion of *trans* junctions for the indicated CIM position used. **(b)** Correlations between predicted and actual efficiencies between tested *ITGB1* TSMs (with dCasRx) with qPCR (Pearson’s correlation *r* = .42, p = .019) and ddPCR (Pearson’s correlation *r* = .69 p = 1.8E-05) readouts. **(c)** Correlation between qPCR and ddPCR readout for the tested *ITGB1* TSMs (Pearson’s correlation *r* = .86, p = 3.8E-10) **(d)** Correlations between predicted and actual efficiencies between *ITGB1* CIMs (with the +231 TSM and dCasRx) with qPCR (Pearson’s correlation *r* = .62, p = 5.9E-05) and ddPCR (Pearson’s correlation *r* = .64, p = 2.9E-04) readouts. **(e)** Correlation between qPCR and ddPCR readout for the tested *ITGB1* TSMs (Pearson’s correlation *r* = .97, p = 2.0E-16)

**Supplementary Figure 5.**
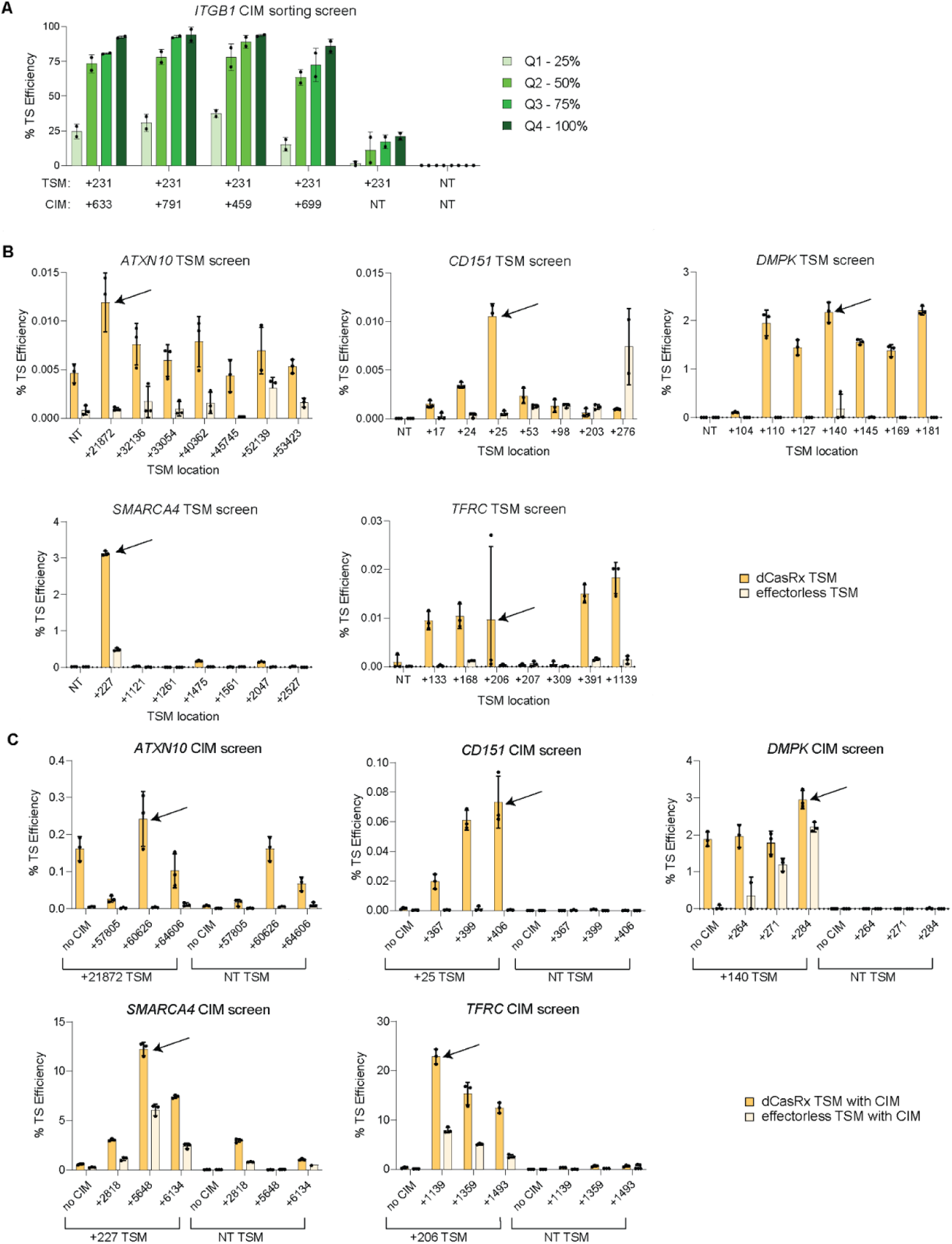
TSM and CIM Spacer Panel for Endogenous Genes. **(a)** Trans-splicing efficiency across quartiles of Cas7-11 expression gated for dCasRx expression with a TSM targeting 231 nucleotides downstream of the splice donor of intron 7 in ITGB1 in conjunction with various CIMs targeting downstream of the splice donor. Data are mean ± s.d.; n = 2 biological replicates, with individual points plotted. **(b)** Comparison of % *trans*-splicing efficiency of each tested TSM for each listed gene with a qPCR readout. Green bars represent conditions with dCasRx as the TSM effector. Gray bars represent conditions with no TSM effector. Arrows indicate which TSM location was used in later experiments. Data are mean ± s.d.; n = 3 biological replicates, with individual points plotted. **(c)** Comparison of % *trans*-splicing efficiency of each tested CIM in conjunction with the listed TSM for each gene with a qPCR readout. Green bars represent conditions with dCasRx as the TSM effector and Cas7-11 as the CIM effector. Gray bars represent conditions with no TSM effector but contain the Cas7-11 CIM effector. Arrows indicate the best performing TSM and CIM combination. Data are mean ± s.d.; n = 3 biological replicates, with individual points plotted.

**Supplementary Figure 6.**
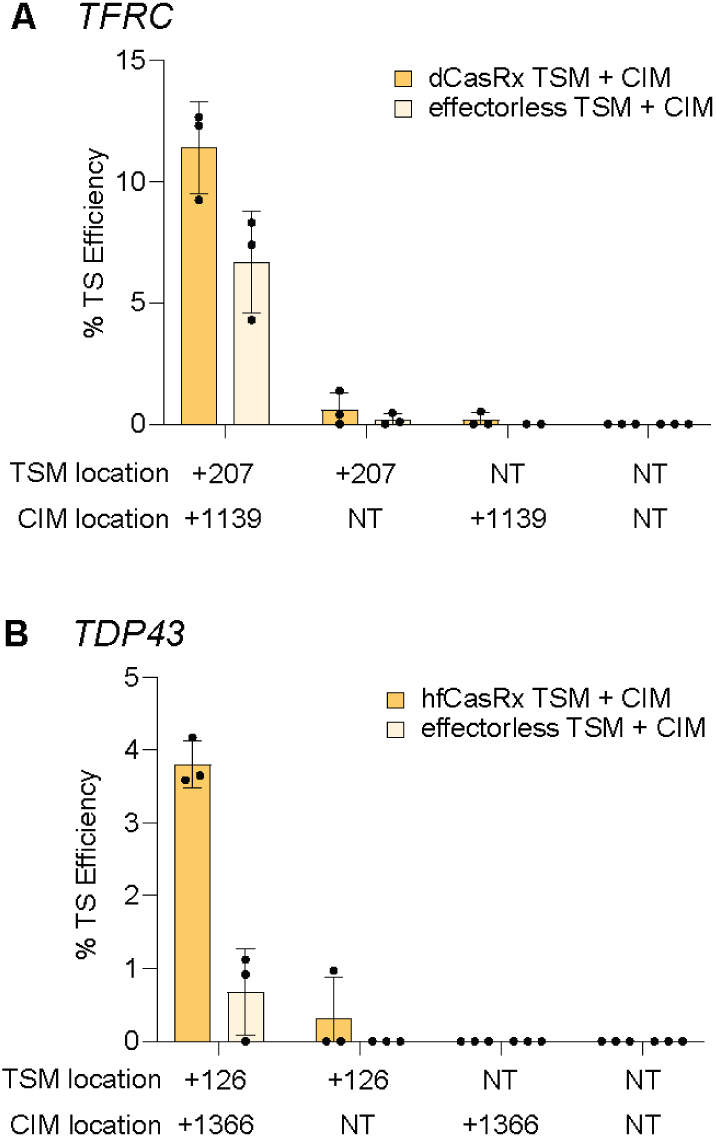
Trans-splicing Endogenous Genes with Non-Reporter Cargo. **(a)** Comparison of different TSM and CIM spacers of % *trans*-splicing efficiency with a ddPCR readout of *TDP43* with a 711 nt cargo carrying the CDS corresponding to exons 5 and 6. Green bars represent conditions with hfCasRx as a TSM effector and Cas7-11 as a CIM effector. Gray bars represent no TSM effector and Cas7-11 as a CIM effector. **(b)** Comparison of different TSM and CIM spacers of % *trans*-splicing efficiency with a ddPCR readout of *TFRC* with a 2.1kb cargo carrying the CDS corresponding to exons 7-19, an XTEN16 linker, and the complete CDS of msfGFP. Green bars represent conditions with dCasRx as a TSM effector and Cas7-11 as a CIM effector. Gray bars represent no TSM effector and Cas7-11 as a CIM effector. All data are mean ± s.d.; n = 3 biological replicates, with individual points plotted.

**Supplementary Figure 7.**
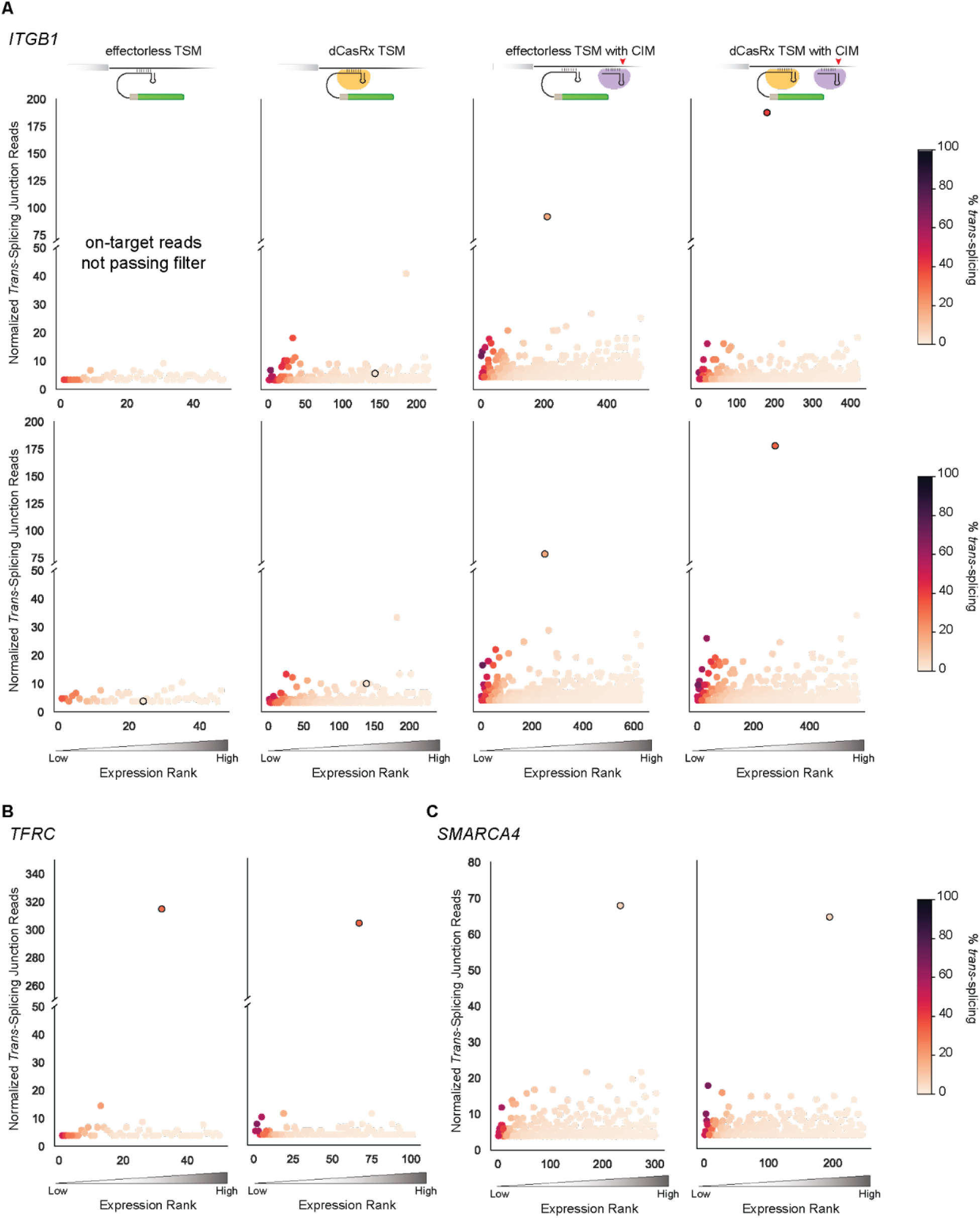
Trans-Splicing Specificity for Endogenous Transcripts. **(a)** A snapshot of trans-splicing efficiency and specificity across all detected *trans*-splicing junctions that passed filtering for different effector combinations targeting *ITGB1*. The first row corresponds to the first RNA-seq replicate and the second row corresponds to the second RNA-seq replicate of each condition, which is separated by column. The effector combination used is represented by the schematic at the top of each column. The rightmost plot of the first row is replicated from Figure 3G for comparison. **(b)** A snapshot of trans-splicing efficiency and specificity across all detected *trans*-splicing junctions that passed filtering for the best TSM and best CIM combination for *SMARCA4* (two rightmost plots) and *TFRC* (two leftmost plots). All plots represent one RNA-seq replicate where each dot corresponds to a *trans*-splicing junction that passed filtering. The x-axis position of each dot represents its expression rank compared to the other *trans*-splicing junctions. The y-axis position of each dot represents the number of normalized *trans*-splicing junctions detected for that junction. The color of each dot represents the *trans*-splicing efficiency of that junction (see legend). The outlined dot represents the on-target *trans*-splicing junction.

**Supplementary Figure 8.**
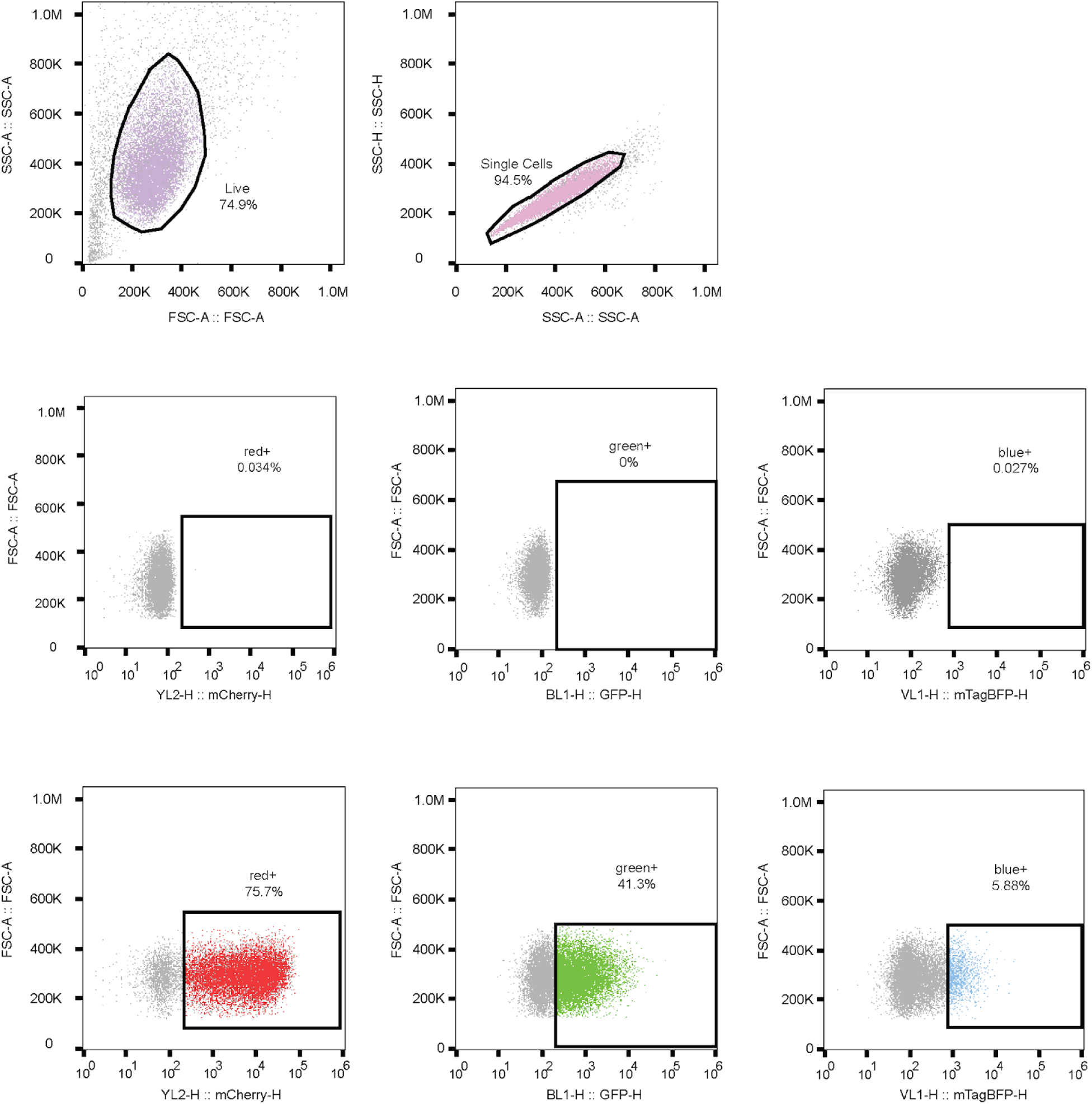
Gating for Flow Cytometry. Gates for live cells were drawn from area of forward scatter and area of side scatter plots, and single cell gates from area side scatter and height of side scatter plots. All mCherry, GFP, and BFP gates were drawn from the heights of the corresponding channels against the area of the forward scatter, and all were gated on single cells, unless otherwise noted in the text.

## REFERENCES

1. Bacon, B. R., Adams, P. C., Kowdley, K. V., Powell, L. W., Tavill, A. S. & Diseases, A. A. for the S. of L. (2011). Diagnosis and management of hemochromatosis: 2011 Practice Guideline by the American Association for the Study of Liver Diseases. Hepatology, 54(1), 328–343. 10.1002/hep.24330

2. Balendra, R. & Isaacs, A. M. (2018). C9orf72-mediated ALS and FTD: multiple pathways to disease. Nature Reviews Neurology, 14(9), 544–558. 10.1038/s41582-018-0047-2

3. Borrajo, J., Javanmardi, K., Griffin, J., Martin, S. J. S., Yao, D., Hill, K., Blainey, P. & Al-Shayeb, B. (2023). Programmable multi-kilobase RNA editing using CRISPR-mediated trans-splicing. bioRxiv. 10.1101/2023.08.18.553620

4. Buratti, E. (2015). Chapter One Functional Significance of TDP-43 Mutations in Disease. Advances in Genetics, 91, 1–53. 10.1016/bs.adgen.2015.07.001

5. Coady, T. H., Baughan, T. D., Shababi, M., Passini, M. A. & Lorson, C. L. (2008). Development of a Single Vector System that Enhances Trans-Splicing of SMN2 Transcripts. PLoS ONE, 3(10), e3468. 10.1371/journal.pone.0003468

6. Depienne, C. & Mandel, J.-L. (2021). 30 years of repeat expansion disorders: What have we learned and what are the remaining challenges? The American Journal of Human Genetics, 108(5), 764–785. 10.1016/j.ajhg.2021.03.011

7. Dobin, A., Davis, C. A., Schlesinger, F., Drenkow, J., Zaleski, C., Jha, S., Batut, P., Chaisson, M. & Gingeras, T. R. (2013). STAR: ultrafast universal RNA-seq aligner. Bioinformatics, 29(1), 15–21. 10.1093/bioinformatics/bts635

8. Drumm, M. L., Ziady, A. G. & Davis, P. B. (2012). Genetic Variation and Clinical Heterogeneity in Cystic Fibrosis. Annual Review of Pathology: Mechanisms of Disease, 7(1), 267–282. 10.1146/annurev-pathol-011811-120900

9. Du, M., Jillette, N., Zhu, J. J., Li, S. & Cheng, A. W. (2020). CRISPR artificial splicing factors. Nature Communications, 11(1), 2973. 10.1038/s41467-020-16806-4

10. Emig, D., Salomonis, N., Baumbach, J., Lengauer, T., Conklin, B. R. & Albrecht, M. (2010). AltAnalyze and DomainGraph: analyzing and visualizing exon expression data. Nucleic Acids Research, 38(suppl_2), W755–W762. 10.1093/nar/gkq405

11. Gijselinck, I., Cruts, M. & Broeckhoven, C. V. (2018). The Genetics of C9orf72 Expansions. Cold Spring Harbor Perspectives in Medicine, 8(4), a026757. 10.1101/cshperspect.a026757

12. Han, S., Zhao, B. S., Myers, S. A., Carr, S. A., He, C. & Ting, A. Y. (2020). RNA–protein interaction mapping via MS2- or Cas*13*-based APEX targeting. Proceedings of the National Academy of Sciences, 117(36), 22068–22079. 10.1073/pnas.2006617117

13. Hanany, M., Rivolta, C. & Sharon, D. (2020). Worldwide carrier frequency and genetic prevalence of autosomal recessive inherited retinal diseases. Proceedings of the National Academy of Sciences, 117(5), 2710–2716. 10.1073/pnas.1913179117

14. Joung, J., Konermann, S., Gootenberg, J. S., Abudayyeh, O. O., Platt, R. J., Brigham, M. D., Sanjana, N. E. & Zhang, F. (2017). Genome-scale CRISPR-Cas9 knockout and transcriptional activation screening. Nature Protocols, 12(4), 828–863. 10.1038/nprot.2017.016

15. Kaseniit, K. E., Katz, N., Kolber, N. S., Call, C. C., Wengier, D. L., Cody, W. B., Sattely, E. S. & Gao, X. J. (2022). Modular, programmable RNA sensing using ADAR editing in living cells. Nature Biotechnology, 1–6. 10.1038/s41587-022-01493-x

16. Kawakami, M., Ishikawa, R., Amano, Y., Sunohara, M., Watanabe, K., Ohishi, N., Yatomi, Y., Nakajima, J., Fukayama, M., Nagase, T. & Takai, D. (2013). Detection of novel paraja ring finger 2-fer tyrosine kinase mRNA chimeras is associated with poor postoperative prognosis in non-small cell lung cancer. Cancer Science, 104(11), 1447–1454. 10.1111/cas.12250

17. Kikumori, T., Cote, G. J. & Gagel, R. F. (2001). Promiscuity of Pre-mRNA Spliceosome-Mediated Trans Splicing: A Problem for *Gene Therapy*? Human Gene Therapy, 12(11), 1429–1441. 10.1089/104303401750298580

18. Konermann, S., Lotfy, P., Brideau, N. J., Oki, J., Shokhirev, M. N. & Hsu, P. D. (2018). Transcriptome Engineering with RNA-Targeting Type VI-D CRISPR Effectors. Cell, 173(3), 665–676.e14. 10.1016/j.cell.2018.02.033

19. Krause, M. & Hirsh, D. (1987). A trans-spliced leader sequence on actin mRNA in C. elegans. Cell, 49(6), 753–761. 10.1016/0092-8674(87)90613-1

20. Li, H., Wang, J., Mor, G. & Sklar, J. (2008). A Neoplastic Gene Fusion Mimics Trans-Splicing of RNAs in Normal Human Cells. Science, 321(5894), 1357–1361. 10.1126/science.1156725

21. Liemberger, B., Hofbauer, J. P., Wally, V., Arzt, C., Hainzl, S., Kocher, T., Murauer, E. M., Bauer, J. W., Reichelt, J. & Koller, U. (2018). RNA Trans-Splicing Modulation via Antisense Molecule Interference. International Journal of Molecular Sciences, 19(3), 762. 10.3390/ijms19030762

22. Liu, X., Jiang, Q., Mansfield, S. G., Puttaraju, M., Zhang, Y., Zhou, W., Cohn, J. A., Garcia-Blanco, M. A., Mitchell, L. G. & Engelhardt, J. F. (2002). Partial correction of endogenous ΔF508 CFTR in human cystic fibrosis airway epithelia by spliceosome-mediated RNA trans-splicing. Nature Biotechnology, 20(1), 47–52. 10.1038/nbt0102-47

23. Mansfield, S. G., Kole, J., Puttaraju, M., Yang, C. C., Garcia-Blanco, M. A., Cohn, J. A. & Mitchell, L. G. (2000). Repair of CFTR mRNA by spliceosome-mediated RNA trans-splicing. Gene Therapy, 7(22), 1885–1895. 10.1038/sj.gt.3301307

24. Marasco, L. E. & Kornblihtt, A. R. (2023). The physiology of alternative splicing. Nature Reviews Molecular Cell Biology, 24(4), 242–254. 10.1038/s41580-022-00545-z

25. Matera, A. G. & Wang, Z. (2014). A day in the life of the spliceosome. Nature Reviews Molecular Cell Biology, 15(2), 108–121. 10.1038/nrm3742

26. Murauer, E. M., Gache, Y., Gratz, I. K., Klausegger, A., Muss, W., Gruber, C., Meneguzzi, G., Hintner, H. & Bauer, J. W. (2011). Functional Correction of Type VII Collagen Expression in Dystrophic Epidermolysis Bullosa. Journal of Investigative Dermatology, 131(1), 74–83. 10.1038/jid.2010.249

27. Nikom, D. & Zheng, S. (2023). Alternative splicing in neurodegenerative disease and the promise of RNA therapies. Nature Reviews Neuroscience, 24(8), 457–473. 10.1038/s41583-023-00717-6

28. Nuñez, J. K., Chen, J., Pommier, G. C., Cogan, J. Z., Replogle, J. M., Adriaens, C., Ramadoss, G. N., Shi, Q., Hung, K. L., Samelson, A. J., Pogson, A. N., Kim, J. Y. S., Chung, A., Leonetti, M. D., Chang, H. Y., Kampmann, M., Bernstein, B. E., Hovestadt, V., Gilbert, L. A. & Weissman, J. S. (2021). Genome-wide programmable transcriptional memory by CRISPR-based epigenome editing. Cell, 184(9), 2503–2519.e17. 10.1016/j.cell.2021.03.025

29. Orengo, J. P., Bundman, D. & Cooper, T. A. (2006). A bichromatic fluorescent reporter for cell-based screens of alternative splicing. Nucleic Acids Research, 34(22), e148–e148. 10.1093/nar/gkl967

30. Özcan, A., Krajeski, R., Ioannidi, E., Lee, B., Gardner, A., Makarova, K. S., Koonin, E. V., Abudayyeh, O. O. & Gootenberg, J. S. (2021). Programmable RNA targeting with the single-protein CRISPR effector Cas7-11. Nature, 597(7878), 720–725. 10.1038/s41586-021-03886-5

31. Pickrell, J. K., Pai, A. A., Gilad, Y. & Pritchard, J. K. (2010). Noisy Splicing Drives mRNA Isoform Diversity in Human Cells. PLoS Genetics, 6(12), e1001236. 10.1371/journal.pgen.1001236

32. Pistoni, M., Ghigna, C. & Gabellini, D. (2010). Alternative splicing and muscular dystrophy. RNA Biology, 7(4), 441–452. 10.4161/rna.7.4.12258

33. Ploeg, L. H. T. V. der, Liu, A. Y. C., Michels, P. A. M., Lange, T. D., Borst, P., Majumder, H. K., Weber, H., Veeneman, G. H. & Boom, J. V. (1982). RNA splicing is required to make the messenger RNA for a variant surface antigen in trypanosomes. Nucleic Acids Research, 10(12), 3591–3604. 10.1093/nar/10.12.3591

34. Pramono, Z. A. D., Takeshima, Y., Alimsardjono, H., Ishii, A., Takeda, S. & Matsuo, M. (1996). Induction of Exon Skipping of the Dystrophin Transcript in Lymphoblastoid Cells by Transfecting an Antisense Oligodeoxynucleotide Complementary to an Exon Recognition Sequence. Biochemical and Biophysical Research Communications, 226(2), 445–449. 10.1006/bbrc.1996.1375

35. Puttaraju, M., Jamison, S. F., Mansfield, S. G., Garcia-Blanco, M. A. & Mitchell, L. G. (1999). Spliceosome-mediated RNA trans-splicing as a tool for gene therapy. Nature Biotechnology, 17(3), 246–252. 10.1038/6986

36. Roca, X., Krainer, A. R. & Eperon, I. C. (2013). Pick one, but be quick: 5′ splice sites and the problems of too many choices. Genes & Development, 27(2), 129–144. 10.1101/gad.209759.112

37. Rogalska, M. E., Vivori, C. & Valcárcel, J. (2023). Regulation of pre-mRNA splicing: roles in physiology and disease, and therapeutic prospects. Nature Reviews Genetics, 24(4), 251–269. 10.1038/s41576-022-00556-8

38. Rovai, A., Chung, B., Hu, Q., Hook, S., Yuan, Q., Kempf, T., Schmidt, F., Grimm, D., Talbot, S. R., Steinbrück, L., Götting, J., Bohne, J., Krooss, S. A. & Ott, M. (2022). In vivo adenine base editing reverts C282Y and improves iron metabolism in hemochromatosis mice. Nature Communications, 13(1), 5215. 10.1038/s41467-022-32906-9

39. Scotti, M. M. & Swanson, M. S. (2016). RNA mis-splicing in disease. Nature Reviews Genetics, 17(1), 19–32. 10.1038/nrg.2015.3

40. Skandalis, A. (2016). Estimation of the minimum mRNA splicing error rate in vertebrates. Mutation Research/Fundamental and Molecular Mechanisms of Mutagenesis, 784, 34–38. 10.1016/j.mrfmmm.2016.01.002

41. Tockner, B., Kocher, T., Hainzl, S., Reichelt, J., Bauer, J. W., Koller, U. & Murauer, E. M. (2016). Construction and validation of an RNA trans-splicing molecule suitable to repair a large number of COL7A1 mutations. Gene Therapy, 23(11), 775–784. 10.1038/gt.2016.57

42. Tong, H., Huang, J., Xiao, Q., He, B., Dong, X., Liu, Y., Yang, X., Han, D., Wang, Z., Wang, X., Ying, W., Zhang, R., Wei, Y., Xu, C., Zhou, Y., Li, Y., Cai, M., Wang, Q., Xue, M., … Yang, H. (2022). High-fidelity Cas13 variants for targeted RNA degradation with minimal collateral effects. Nature Biotechnology, 1–12. 10.1038/s41587-022-01419-7

43. Uckun, F. M., Qazi, S., Ma, H., Reaman, G. H. & Mitchell, L. G. (2015). CD22ΔE12 as a molecular target for corrective repair using RNA trans-splicing: anti-leukemic activity of a rationally designed RNA trans-splicing molecule. Integrative Biology, 7(2), 237–249. 10.1039/c4ib00221k

44. Vecchi, C., Montosi, G. & Pietrangelo, A. (2010). Huh-7: A human “hemochromatotic” cell line. Hepatology, 51(2), 654–659. 10.1002/hep.23410

45. Viles, K. D. & Sullenger, B. A. (2008). Proximity-dependent and proximity-independent trans-splicing in mammalian cells. RNA, 14(6), 1081–1094. 10.1261/rna.384808

46. Wallace, D. F. & Subramaniam, V. N. (2016). The global prevalence of HFE and non-HFE hemochromatosis estimated from analysis of next-generation sequencing data. Genetics in Medicine, 18(6), 618–626. 10.1038/gim.2015.140

47. Wally, V., Murauer, E. M. & Bauer, J. W. (2012). Spliceosome-Mediated Trans-Splicing: The Therapeutic Cut and Paste. Journal of Investigative Dermatology, 132(8), 1959–1966. 10.1038/jid.2012.101

48. Wei, J., Lotfy, P., Faizi, K., Baungaard, S., Gibson, E., Wang, E., Slabodkin, H., Kinnaman, E., Chandrasekaran, S., Kitano, H., Durrant, M. G., Duffy, C. V., Pawluk, A., Hsu, P. D. & Konermann, S. (2023). Deep learning and CRISPR-Cas13d ortholog discovery for optimized RNA targeting. Cell Systems. 10.1016/j.cels.2023.11.006

49. Wolin, S. L. & Maquat, L. E. (2019). Cellular RNA surveillance in health and disease. Science, 366(6467), 822–827. 10.1126/science.aax2957

50. Ye, J., Coulouris, G., Zaretskaya, I., Cutcutache, I., Rozen, S. & Madden, T. L. (2012). Primer-BLAST: A tool to design target-specific primers for polymerase chain reaction. BMC Bioinformatics, 13(1), 134. 10.1186/1471-2105-13-134

51. Zeballos C., M. A., Moore, H. J., Smith, T. J., Powell, J. E., Ahsan, N. S., Zhang, S. & Gaj, T. (2023). Mitigating a TDP-43 proteinopathy by targeting ataxin-2 using RNA-targeting CRISPR effector proteins. Nature Communications, 14(1), 6492. 10.1038/s41467-023-42147-z

